# Expanding the scope of bacterial CRISPR activation with PAM-flexible dCas9 variants

**DOI:** 10.1101/2022.07.14.500123

**Authors:** Cholpisit Kiattisewee, Ava V. Karanjia, Mateusz Legut, Zharko Daniloski, Samantha E. Koplik, Joely Nelson, Benjamin P. Kleinstiver, Neville E. Sanjana, James M. Carothers, Jesse G. Zalatan

**Affiliations:** Molecular Engineering & Sciences Institute and Center for Synthetic Biology, University of Washington, Seattle, WA 98195, United States; Department of Chemical Engineering, University of Washington, Seattle, WA 98195, United States; New York Genome Center, New York, NY 10013, United States; Department of Biology New York, University New York, NY 10013, United States; Department of Bioengineering, University of Washington, Seattle, WA 98195, United States; Center for Genomic Medicine, Massachusetts General Hospital, Boston, MA 02114, United States; Department of Pathology, Massachusetts General Hospital, Boston, MA 02114, United States; Department of Pathology, Harvard Medical School, Boston, MA 02115, United States; Department of Chemistry, University of Washington, Seattle, WA 98195, United States

## Abstract

CRISPR-Cas transcriptional tools have been widely applied for programmable regulation of complex biological networks. In comparison to eukaryotic systems, bacterial CRISPR activation (CRISPRa) has stringent target site requirements for effective gene activation. While genes may not always have an NGG PAM at the appropriate position, PAM-flexible dCas9 variants can expand the range of targetable sites. Here we systematically evaluate a panel of PAM-flexible dCas9 variants for their ability to activate bacterial genes. We observe that dxCas9-NG provides a high dynamic range of gene activation for sites with NGN PAMs while dSpRY permits modest activity across almost any PAM. Similar trends were observed for heterologous and endogenous promoters. For all variants tested, improved PAM-flexibility comes with the tradeoff that CRISPRi-mediated gene repression becomes less effective. Weaker CRISPRi gene repression can be partially rescued by expressing multiple sgRNAs to target many sites in the gene of interest. Our work provides a framework to choose the most effective dCas9 variant for a given set of gene targets, which will further expand the utility of CRISPRa/i gene regulation in bacterial systems.

## 1. Introduction

CRISPR-Cas transcriptional regulation enables programmable control over gene activation and repression in bacteria (Bikard et al., 2013; Dong et al., 2018; Qi et al., 2013). These tools can be used to regulate endogenous gene networks both to probe biological function and to engineer new behaviors. However, effective CRISPR activation (CRISPRa) is limited by complex and stringent target site requirements (Fontana et al., 2020a; Ho et al., 2020; Liu et al., 2019; Villegas Kcam et al., 2021). Previous work has demonstrated a sharply periodic pattern of effective target sites occurring every 10 bp within a 60-100 bp region upstream of the TSS (Dong et al., 2018; Fontana et al., 2020a, 2020b; Kiattisewee et al., 2021). Targeting these sites with *S. pyogenes* dCas9 requires a compatible protospacer adjacent motif (PAM) at the right position, and endogenous genes that lack an appropriate PAM may be incompatible for activation via Sp-Cas9.

We previously demonstrated that one of the first expanded PAM dCas9 variants, dxCas9(3.7) (Hu et al., 2018), could activate some genes that lacked an appropriately-positioned NGG PAM (Fontana et al., 2020a). Specifically, dxCas9(3.7) produced at least two-fold increases in gene expression at three out of seven endogenous promoters tested (Fontana et al., 2020a). Although these results were encouraging, all seven of the candidate promoters were predicted to encode PAMs compatible with dxCas9(3.7) and all seven were expected to be activated. Since our initial work with dxCas9(3.7), several new expanded PAM dCas9 variants have been described (Legut et al., 2020; Nishimasu et al., 2018; Walton et al., 2020), raising the possibility that improved variants could further expand the number of targetable genes for CRISPRa in bacteria.

In this work, we demonstrate that PAM-flexible dCas9 variants can improve transcriptional activation at endogenous genes compared to both dCas9 and dxCas9(3.7). We observe activation at previously inaccessible gene targets, and we observe a tradeoff between fold-activation and PAM flexibility. We also demonstrate that expanded PAM dCas9 variants are partially impaired for CRISPR interference (CRISPRi) gene repression. This effect can be mitigated by targeting multiple CRISPR complexes to the desired gene. By systematically characterizing the properties of PAM-flexible dCas9 variants in bacterial CRISPRa/i, we provide a framework to choose the most effective variant for a given gene target or set of targets. This toolbox of dCas9 variants will further expand the utility of CRISPRa/i in bacterial systems for a broad range of applications including metabolic engineering and genome-wide functional screens.

## 2. Results

### 2.1 *In-silico* PAM availability analysis to predict effective CRISPRa sites

In bacterial systems, effective CRISPRa requires a target site that is precisely positioned upstream of the gene of interest (Fontana et al., 2020a). Consequently, the ability to activate an arbitrary gene is dependent on an appropriately positioned PAM at the desired target site. The widely-used *S. pyogenes* Cas9 (Sp-Cas9) is generally limited to NGG PAMs, but several engineered Cas9 variants have been developed with more flexibility to accommodate various PAMs. We examined two groups of engineered Cas9 variants (Figure 1A). The first group includes xCas9-NG, a variant that exhibits high nuclease efficiency and CRISPRa/i performance at NGN PAMs in mammalian systems (Hu et al., 2018; Legut et al., 2020; Nishimasu et al., 2018). The second group includes SpG and SpRY (Kleinstiver et al., 2016, 2015; Nishimasu et al., 2018; Walton et al., 2021, 2020). The SpRY variant was engineered to be near-PAMless with some preference for NRN PAMs.

**Figure 1:**
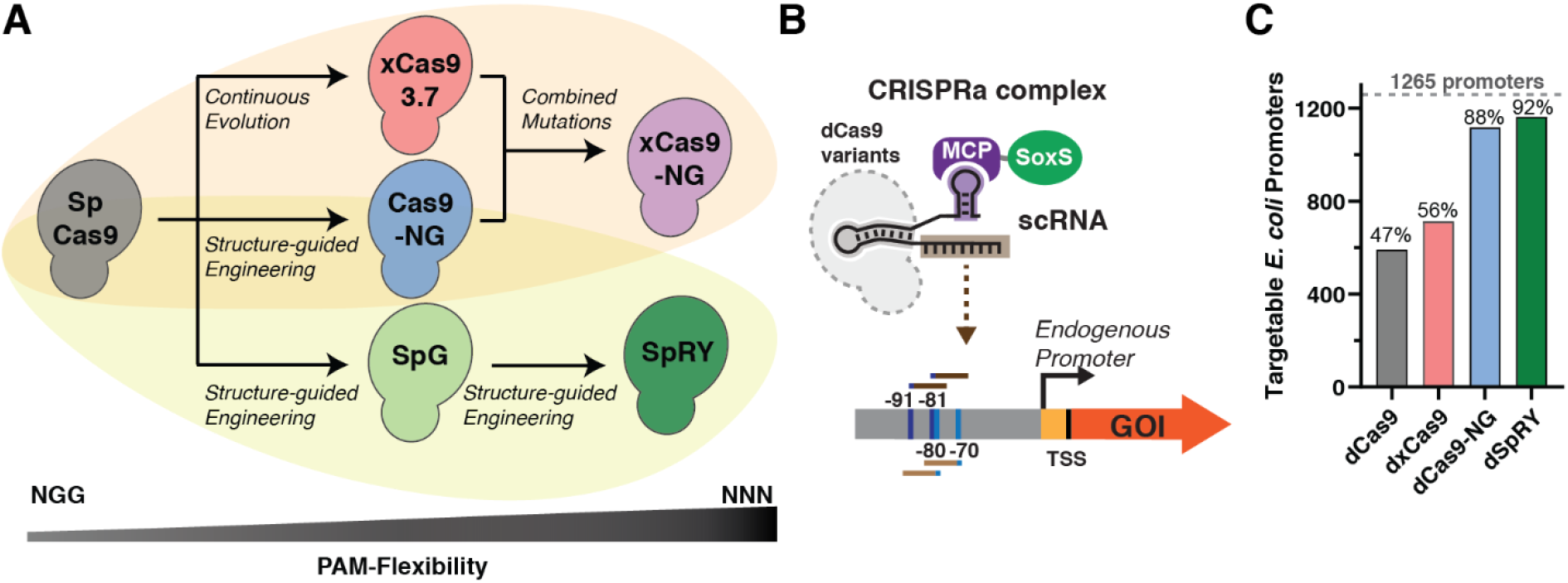
Engineered PAM-flexible Cas9 variants and PAM availability analysis. (A) PAM-flexible S. pyogenes Cas9 (Sp-Cas9) have been engineered using various rational design, screening, and directed evolution methodologies (e.g. phage-assisted continuous evolution (PACE), structure-guided engineering monitored via HT-PAMDA) (Hu et al., 2018; Kleinstiver et al., 2016, 2015; Legut et al., 2020; Nishimasu et al., 2018; Walton et al., 2021, 2020). (B) The CRISPRa complex consists of dCas9, an scRNA, and the MCP-SoxS activator (Dong et al., 2018). Previous work suggests that four precisely-positioned sites upstream of the TSS are most effective for CRISPRa (-70 and -80 for the template strand and -81 and -91 for non-template strands relative to the TSS) (Fontana et al., 2020a). (C) 1265 *E. coli* endogenous promoters were analyzed for target site availability with different PAM-flexible dCas9 variants. Promoters were identified as targetable if they have at least one effective target site with a compatible PAM. For each variant, PAM preferences were obtained from previously-reported nuclease screening experiments (see Methods; PAMs with at least 20% of Sp-Cas9 activity at NGG PAMs were considered compatible).

We previously identified a set of four precisely-positioned sites upstream of the TSS that are the most effective for CRISPRa with the SoxS activator (Figure 1B) (Fontana et al., 2020a). We performed an *in-silico* analysis to identify promoters with accessible PAMs at one or more of the optimal upstream positions in *Escherichia coli* and *Pseudomonas putida*. We restricted the search to promoters with high-confidence TSS positions (Santos-Zavaleta et al., 2019) and with sigma factors previously identified to be effective for CRISPRa with the SoxS activator (Fontana et al., 2020a). Together these criteria were met for 1265 out of 4042 promoters in *E. coli*. PAM compatibility for each Cas9 variant was predicted based on the reported nuclease activity with variable PAM targets (See Methods and Supplementary Methods) (Kim et al., 2020; Legut et al., 2020; Walton et al., 2020). We performed the analysis with 4 deactivated Cas9 variants (dCas9, dxCas9(3.7), dCas9-NG, and dSpRY) of different PAM-flexibility (Table S4). Because xCas9-NG and Cas9-NG exhibit comparable levels of PAM flexibility in mammalian cell assays for nuclease activity and CRISPRa (Figure S2), we expect the predictions for dCas9-NG to be representative for dxCas9-NG. The number of promoters with at least one PAM at a suitable position was 47% for dCas9, and increased to almost all promoters for the engineered variants (89% and 93% for dCas9-NG and dSpRY, respectively) (Figure 1C). A similar trend was observed in *P. putida* (Figure S3). It is important to note that these predictions are based on nuclease activity in mammalian cells and may not accurately predict bacterial CRISPRa activity, since binding and cleavage determinants may differ.

### 2.2 CRISPRa on non-NGG PAM is improved with engineered dCas9 variants

To test whether PAM-flexible Cas9 variants can increase the number of available CRISPRa target sites, we constructed expression cassettes for the catalytically-inactive versions of each Cas9 variant (dCas9, dxCas9(3.7), dCas9-NG, dxCas9-NG, dSpG, and dSpRY). Our bacterial CRISPRa system uses dCas9 and a modified guide RNA (termed scaffold RNA, scRNA) with an MS2 hairpin to recruit the MCP-SoxS activator (Dong et al., 2018; Fontana et al., 2020a) (Figure 2A). We first determined whether each dCas9 variant was effective for CRISPRa at a canonical AGG PAM target site. We used a previously-described reporter gene (J3-BBa_J23117-mRFP) with an AGG PAM positioned at -81 relative to the TSS (Fontana et al., 2020). All tested dCas9 variants exhibited similar activation (∼40-50-fold) with the canonical AGG PAM (Figure 2D).

**Figure 2:**
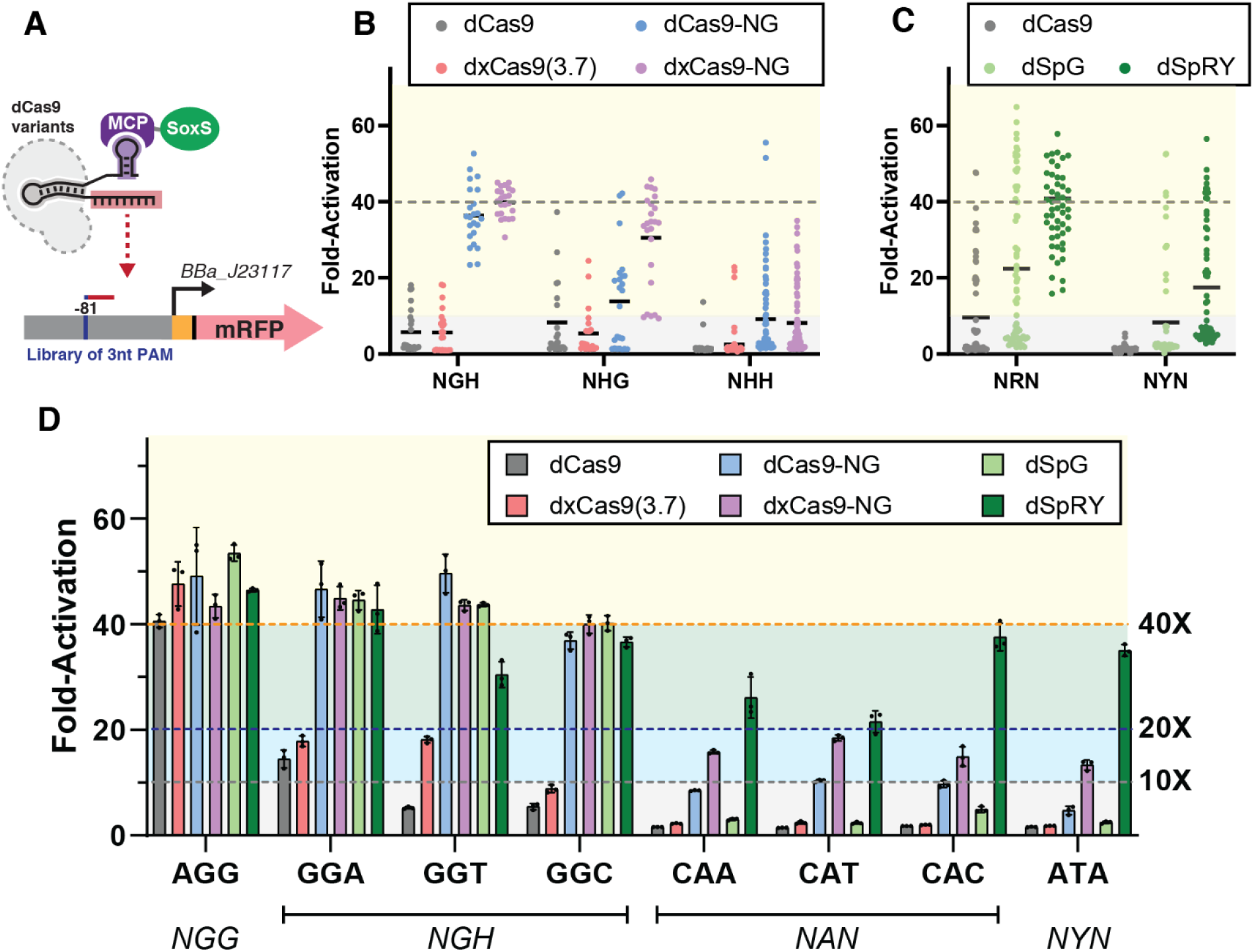
CRISPRa with PAM-flexible dCas9 variants. (A) CRISPRa for PAM-flexible dCas9 variants was tested on an mRFP reporter gene with libraries of alternative 3 nucleotide PAMs at the -81 target site. (B) Reporter gene expression with the dxCas9-NG group (Figure 1A) at NGH, NHG, and NHH PAM libraries. dxCas9-NG outperforms other variants in every PAM library. (C) Reporter gene expression with the dSpRY group (Figure 1A) at NRN and NYN PAM libraries. dSpRY exhibited the highest CRISPRa efficiency. (D) Direct comparisons of reporter gene expression with dCas9, dxCas9(3.7), dCas9-NG, dxCas9-NG, dSpG, and dSpRY at a representative set of PAMs. dCas9-NG, dxCas9-NG, and dSpRY performed best at NGN PAMs while dSpRY outperformed other variants at NAN and NYN PAMs. An scRNA targeting the J306 sequence was used for all library screens and individual PAM assays. To calculate fold-activation, we used an off-target scRNA (hAAVS1) with an AGG PAM reporter to define the basal expression level. Basal reporter expression levels vary <1.5-fold with different dCas9 variants or PAMs (Figure S4). Values in panel D represent the mean ± standard deviation calculated from n = 3.

To evaluate the effectiveness of each dCas9 variant for recognizing non-canonical PAM sites (non-NGG), we constructed libraries of reporters with varied PAM sequences. The first group of dCas9 variants (dxCas9(3.7), dCas9-NG, and dxCas9-NG) has a preference for NGN PAMs, so we screened three PAM libraries (NGH, NHG, and NHH, where H is not G). This approach follows the strategy previously used to characterize several Cas9 variants in mammalian systems (Legut et al., 2020). Taken together, the data indicate that dxCas9-NG provided the highest fold-activation across the largest number of alternative PAM reporters (Figure 2B). dxCas9-NG was effective at all NGH reporters (23 out of 23, or 100% had >10-fold activation) and most NHG reporters (21 out of 24, or 88% had >10-fold activation), but displayed substantially diminished effectiveness at NHH reporters (19 out of 71, or 27% had >10-fold activation). dCas9-NG performed similarly to dxCas9-NG at NGH and NHH PAMs, but notably weaker at NHG PAMs (NGH: 23 out of 23, or 100% had >10-fold activation; NHH: 22 out of 72, or 31% had >10-fold activation; NHG: 13 out of 24, or 54% had >10-fold activation). dCas9 and dxCas9(3.7) led to consistently reduced fold-activation and activated fewer reporters than dxCas9-NG across all three PAM libraries. The broad effectiveness of dxCas9-NG at NGH/NHG PAMs, along with its ability to activate some NHH PAMs, is consistent with prior reports from mammalian systems (Legut et al., 2020).

To evaluate the effectiveness of the dSpRY group of dCas9 variants (Figure 1A), we screened NRN and NYN PAM libraries (R = A or G; Y = C or T) following the previously-determined PAM preferences for dSpRY in mammalian cells (Walton et al., 2021, 2020). As expected, we observed the strongest performance across the broadest range of PAM reporters with dSpRY. This variant produced >10-fold CRISPRa at all tested reporters in the NRN library and 29 out of 64 (45%) of the reporters in the NYN library. In both libraries, dSpRY consistently outperformed both dSpG and dCas9 (Figure 2C).

We proceeded to directly compare the CRISPRa efficiency of dxCas9-NG and dSpRY at specific non-NGG PAMs. We tested reporters with NGN (GGA, GGT, GGC), NAN (CAA, CAT, CAC), and NYN (ATA) PAMs (Figure 2D). We found that dxCas9-NG and dSpRY outperformed both dCas9 and dxCas9(3.7) at all of the non-NGG PAMs. dxCas9-NG exhibited the highest activity at NGN (≥40-fold activation) and moderately weaker activity at NAN and NYN (between 10-fold to 20-fold activation). Compared to dxCas9-NG, dSpRY produced similar or stronger activation at NAN and NYN PAMs (>20-fold activation). These data indicate that different PAM-flexible variants have distinct patterns of optimal PAMs. Together with the PAM library screens, these results suggest that dCas9-NG, dxCas9-NG or dSpG should be used for NGH, dxCas9-NG should be used for for NHG, and dSpRY should be used for NHH PAMs. For subsequent experiments, we prioritized dxCas9-NG for NGH/NHG PAMs and dSpRY for NHH PAMs.

### 2.3 CRISPRi efficiency is impaired with PAM-flexible dCas9 variants

CRISPRi acts by physically blocking transcription, and effective repression in bacteria can generally be obtained by targeting within the promoter or near the beginning of the gene on the non-template strand (Peters et al., 2016; Qi et al., 2013). CRISPRi is not subject to the same stringent target site requirements as CRISPRa, and there are often many canonical NGG PAMs available in the promoter or at the beginning of the gene. However, because we desire to express a single dCas9 protein to simultaneously target multiple genes for CRISPRa or CRISPRi, it is important to evaluate the performance of expanded PAM variants for CRISPRi. Previous work in bacterial and eukaryotic systems suggests that expanded PAM variants exhibit impaired CRISPRi function (Legut et al., 2020; Wang et al., 2021). We proceeded to evaluate the PAM-flexible variants dxCas9(3.7), dxCas9-NG, and dSpRY for CRISPRi gene repression. We tested multiple distinct sgRNA target sites, with one site in the promoter region and two sites within the ORF, all with NGG PAMs (Figure 3A). At each of these sites, dCas9 produces ∼95-fold repression, while the PAM-flexible variants exhibit varying degrees of impaired repression (Figure 3B and Figure S5). For target sites within the ORF, the PAM-flexible dCas9 variants repress gene expression by 5 to 30-fold; these effects are significant but substantially weaker than the ∼95-fold repression obtained with dCas9 at these sites. For the target site at the promoter, the dxCas9(3.7) performs similarly to dCas9, while dxCas9-NG and dSpRY produce weaker repression effects (28-fold and 14-fold, respectively).

**Figure 3:**
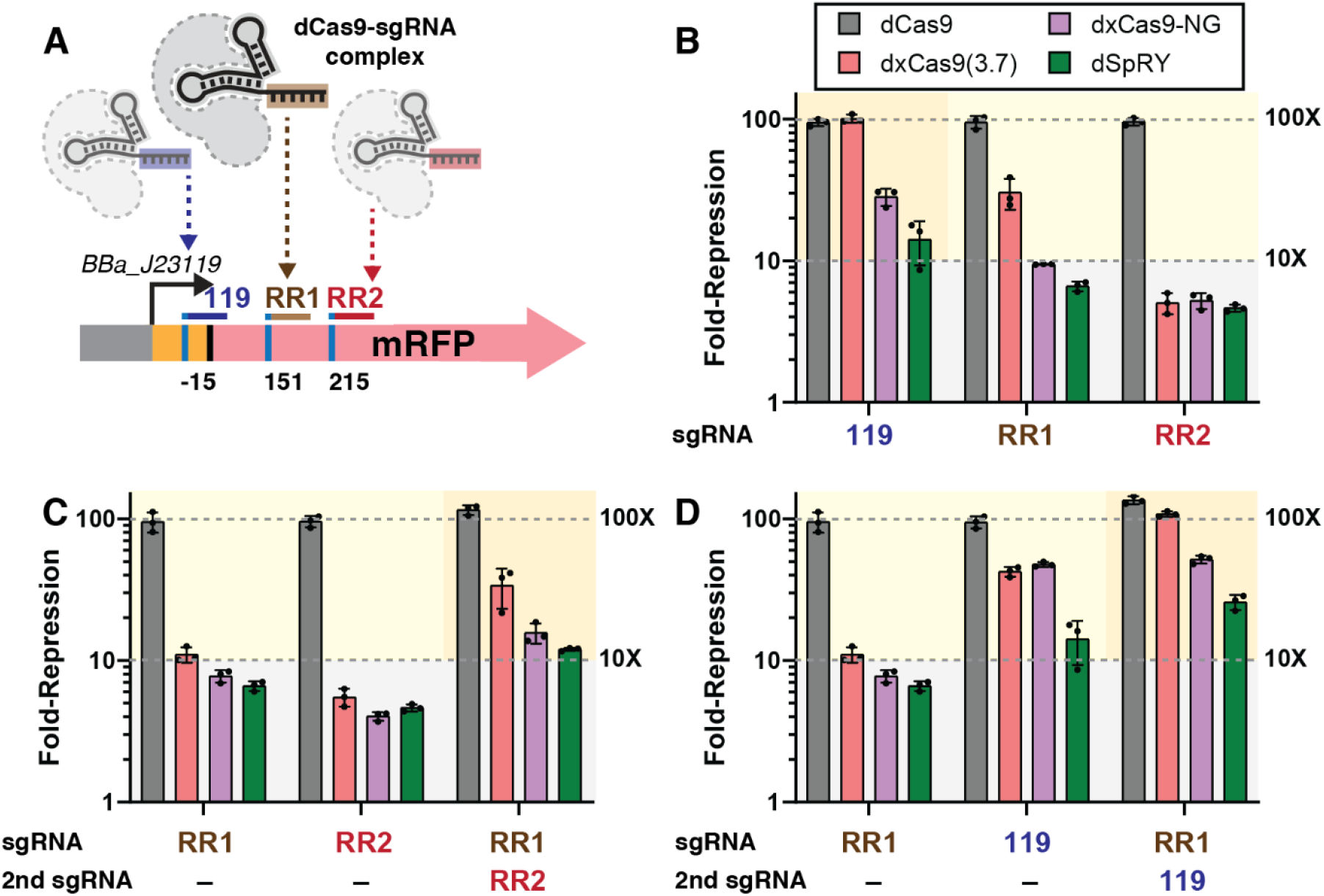
CRISPRi with PAM-flexible dCas9 variants. (A) CRISPRi for PAM-flexible dCas9 variants was tested on an mRFP reporter gene with sgRNAs targeting the promoter (119) or coding sequence (RR1 and RR2), where each target site encodes an NGG PAM. (B) Fold-repression of the mRFP reporter gene with a single expressed sgRNA. (C & D) Comparison of fold-repression with one or two sgRNAs expressed. Fold-repression consistently increases when two sgRNAs are expressed. See Figure S5 for a comparison of all dCas9 variants, including dCas9-NG and dSpG, with single and multiple sgRNAs. Values in panel B, C, and D represent the mean ± standard deviation calculated from n = 3.

CRISPRi repression with dCas9 can be improved by targeting multiple sgRNAs to the same gene (Qi et al., 2013). We therefore tested whether pairs of sgRNAs could be used to improve CRISPRi repression with PAM-flexible dCas9 variants. In each case, we observed improved repression (Figure 3B and 3C). Most notably, dSpRY repression can be improved from 14-fold to 26-fold. The dSpRY variants has one of the broadest targeting ranges due to its wide tolerance of PAMs for CRISPRa (Figure 2D), and the ability to improve its CRISPRi function via gRNA multiplexing suggests that dSpRY can be used for multi-gene CRISPRa/i programs with a large dynamic range of activation and repression.

### 2.4 PAM-flexible dCas9 variants improve CRISPRa at endogenous gene targets

We previously demonstrated that the expanded PAM variant dxCas9(3.7) enables activation of some endogenous genes that are inaccessible to dCas9 (Fontana et al., 2020a). To determine if dxCas9-NG and dSpRY further increase the pool of activatable endogenous genes, we examined four endogenous promoters previously tested with dxCas9(3.7): yajGp, uxuRp, araEp, and ppiDp2. For each endogenous promoter, we tested one NGG and one non-NGG PAM at appropriate positions upstream of the TSS at -70 or -80 (template strand) or at -81 or -91 (non-template strand), following the targeting rules defined previously (Fontana et al., 2020a). For two promoters previously activated by dxCas9(3.7), yajGp and uxuRp, dxCas9-NG and dSpRY produced comparable or improved activation compared to dxCas9(3.7) (Figure 4B). We also observed that two promoters that could not be activated by dxCas9(3.7) were slightly activated with dxCas9-NG and dSpRY: araEp was activated by 1.3-fold by dxCas9-NG while ppiDp2 was activated by 3.1-fold by dSpRY (Figure S6C).

**Figure 4:**
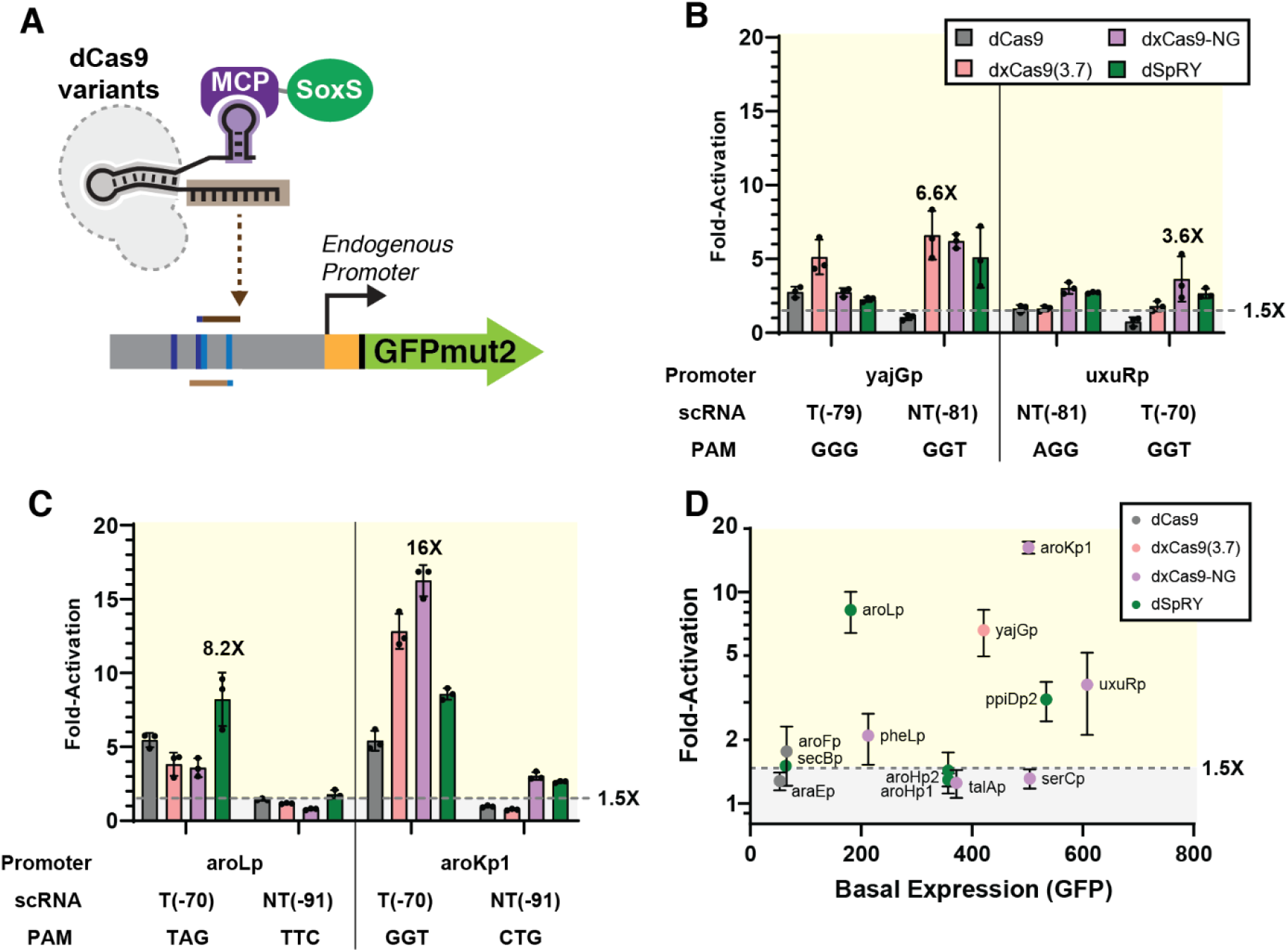
CRISPRa at endogenous promoters is enhanced with PAM-flexible dCas9 variants. (A) CRISPRa at endogenous promoters of *E. coli* was tested using two of the four optimal scRNA positions for each promoter (see Supplemental Methods). (B) CRISPRa at endogenous promoters previously tested with dxCas9(3.7) (yajGp and uxuRp) using NGG or non-NGG PAMs. See Figure S6C for araEp and ppiDp2 promoters. (C) CRISPRa at endogenous promoters involved in aromatic amino acid biosynthesis using non-NGG PAMs. See Figure S6D for additional promoters (D) Plot of fold-activation versus basal expression level for all endogenous promoters tested in (B) & (C). See Table S5 for additional details. Data were collected by flow cytometry and fold-activation was calculated relative to a strain expressing the corresponding dCas9 variant and an off-target hAAVS1 scRNA. Values in panel B, C, and D represent the mean ± standard deviation calculated from n = 3.

We also tested nine new weakly-expressed endogenous promoters chosen from metabolic pathways related to aromatic amino acid biosynthesis (Zaslaver et al., 2006) (See Supplementary Methods). None of these promoters have NGG PAMs at the ideal distances relative to the TSS for CRISPR activation. For each promoter, we identified two candidate non-NGG targets that we expected to be compatible with dxCas9-NG or dSpRY (see Table S3). These sites were selected with the target site position rules described above (Fontana et al., 2020a). Out of nine new promoters tested, five can be activated by more than 1.5-fold. Activation of aroKp1 by dxCas9-NG and activation of aroLp by dSpRY demonstrated the largest fold-activation values (16-fold and 8-fold, respectively) (Figure 4C). We observed modest, ∼2-fold activation for pheLp, secBp, and aroFp, and no activation (<1.5-fold) for aroHp1, aroHp2, talAp, and serCp (Figure S6D). For all promoters that were activated >1.5-fold, we observed distinct behaviors with different PAM-flexible variants. For aroKp1, dxCas9-NG significantly outperforms other variants at the GGT PAM. For aroLp, dSpRY outperforms other variants at the TAG PAM. These behaviors are consistent with our results from the mRFP reporter assay, which suggested the use of dxCas9-NG at NGN PAMs and dSpRY at NHH PAMs (Figure 2D). Taken together, these results suggest that the newer PAM-flexible variants dxCas9-NG and dSpRY can outperform dCas9 and dxCas9(3.7) for bacterial CRISPRa.

Out of the 13 total endogenous promoters tested, we found that five could not be activated above the 1.5-fold threshold (Figure 4B&C, S6C&D). One possible explanation for the failure of some target promoters to activate is that their basal expression levels are too high or too low. We previously observed that bacterial CRISPRa is sensitive to basal promoter strength, with no activation at the weakest promoters, effective activation at moderately weak promoters, and progressively smaller increases in gene expression as basal expression levels increased (Fontana et al., 2020a). Promoters with innately high expression levels may be inaccessible to high activation (>1.5-fold) due to the metabolic burden associated with increasing protein concentration (Gyorgy et al., 2015; Wu et al., 2016). However, none of the five inaccessible promoters have unusually high basal expression levels (Figure 4D). Four of the promoters with <1.5 fold activation (aroHp1, aroHp2, talAp, and serCp) fall in a basal expression range that is higher than aroLp and lower than aroKp1, the two most highly activated promoters. Other factors beyond basal expression levels may be responsible for the failure of these four promoters to activate. The remaining inaccessible promoter, araEp, has the lowest basal expression level of all promoters tested and may be too weak to be activated with our current CRISPRa system.

## 3. Discussion

In this work, we have demonstrated that PAM-flexible dCas9 variants can improve bacterial CRISPRa in both synthetic and endogenous promoter contexts. Target distance from the TSS has been previously shown to be an important factor for effective bacterial CRISPRa (Fontana et al., 2020a; Ho et al., 2020; Liu et al., 2019; Villegas Kcam et al., 2021). PAM-flexible variants expand the scope of accessible PAM sites and therefore enable targeting at precise, optimal positions upstream of the TSS.

Although PAM-flexible dCas9 variants enable CRISPRa at previously inaccessible gene targets, not all endogenous promoters with target sites at the appropriate position were able to be activated (Figure 4). Additional native regulatory machinery at endogenous gene targets may be responsible for preventing activation in these cases. We have previously shown that transcription factor binding can interfere with bacterial CRISPRa, presumably by physically obstructing binding of the CRISPRa complex or blocking the SoxS effector protein from engaging with RNA polymerase (Fontana et al., 2020a). We hypothesize that cryptic and unannotated transcription factor binding sites could be responsible for preventing activation at the endogenous genes targeted with PAM-flexible dCas9 variants in this work.

Improved bacterial CRISPRa with PAM-flexible variants comes with tradeoffs. We found that PAM-flexible dCas9 variants exhibited weaker CRISPRi-based gene repression compared to dCas9 (Figure 3B). One possible explanation for this behavior follows from the observation that increasing PAM promiscuity reduces affinity towards NGG PAMs (Corsi et al., 2022; Hu et al., 2018; Legut et al., 2020; Nishimasu et al., 2018; Walton et al., 2020). This reduced PAM-binding affinity could allow RNA polymerase to more readily displace the CRISPRi complex. Alternatively, a near-PAMless dCas9 variant would be expected to interrogate almost every DNA sequence in the bacterial genome (Anders et al., 2014; Collias and Beisel, 2021) and increase the time needed to find the correct target site. During every cell division, the CRISPRi complex is displaced from the genome (Qi et al., 2013), and increased time will be needed to bind the target site (Jones et al., 2017), which could allow for increased leaky expression and consequently weaker CRISPRi compared to the parent dCas9.

To implement complex, programmable genetic regulatory networks, both upregulation and downregulation controls with broad dynamic ranges are desirable (Alon, 2007; Brophy and Voigt, 2014; Tickman et al., 2021). Thus, identifying systems that improve CRISPRa while maintaining effective CRISPRi is crucial. Multiplexing the CRISPRi targets with additional sgRNAs led to significantly higher fold-repression, although still weaker than CRISPRi with the parent dCas9 (FIgure 3C-3D). It remains to be seen whether these improvements are sufficient for genetic circuit applications. If stronger repression is needed, an alternative approach could be to distribute each engineering task to a different subpool of Cas9 proteins (Collias and Beisel, 2021). In this application, a PAM-flexible variant could be used for CRISPRa and an orthogonal Cas protein could be used for CRISPRi.

These improvements to bacterial CRISPRa bring us closer to the long-standing goal of performing CRISPRa gain-of-function screens for basic discovery and engineering applications. Previously, some successes have been reported for activating natural product biosynthesis pathways (Ameruoso et al., 2022; Ke et al., 2022) but the ability to perform genome-wide CRISPRa screens in bacteria lags far behind eukaryotic systems (Joung et al., 2017; Kampmann, 2018; Sanson et al., 2018; Schmidt et al., 2022). By unlocking these capabilities in bacteria, CRISPRa screens could enable rapid functional annotation of uncharacterized genes. For bioindustrial applications, we envision identifying genes that can overcome bottlenecks in routing metabolic flux or that confer robustness in harsh, non-native growth conditions. In the long term, combined CRISPRi and CRISPRa screens could provide even more information to map biological functions and deconvolute regulatory networks. In mammalian cells, combining information from single gene CRISPRa/i screens enabled improved chemical genetic profiling to identify drug targets (Jost et al., 2017), and dual-gene activation/repression screens have identified genetic interactions and functional relationships between genes (Boettcher et al., 2018; Najm et al., 2018). Combined CRISPRa/i screens should be possible in bacteria, as CRISPRi loss-of-function screens have been broadly applied (Todor et al., 2021), and it is straightforward to target multiple genes with multiple gRNAs for activation and repression (Dong et al., 2018; Kiattisewee et al., 2021; Wu et al., 2020). Some of these goals are plausibly within reach in bacteria using current CRISPRa tools, and further improvements in CRISPRa systems at endogenous gene targets will enable rapid progress in basic research and bioindustrial applications.

## Acknowledgements

We thank Benjamin I. Tickman, Ian D. Faulkner, Diego Alba Burbano, Ryan Cardiff, and members of Carothers and Zalatan laboratories for thorough discussion during the experimental planning and data collection/analysis strategy. We also thank Madelynn N. Whittaker for cloning the original bacterial codon optimized dSpG and dSpRY constructs. This work was supported by NSF Award 1817623 (J.M.C, J.G.Z.), DOE Award DE-EE0008927 (J.M.C, J.G.Z.), and NIH/NHGRI DP2HG010099 (N.E.S.). B.P.K. acknowledges funding from an MGH ECOR Howard M. Goodman Award. This material is based upon work supported by the National Science Foundation Graduate Research Fellowship Program under Grant No. DGE-2140004 (A.V.K.). Any opinions, findings, and conclusions or recommendations expressed in this material are those of the author(s) and do not necessarily reflect the views of the National Science Foundation.

## Author contributions

C.K., A.V.K., J.M.C., and J.G.Z. developed the initial concept; B.P.K. and N.E.S. provided additional insight to the project; C.K. and J.N. performed bioinformatic analysis; C.K., A.V.K., and S.K. performed experiments in bacteria and analyzed the results; M.L. and Z.D. performed experiments in mammalian cells; C.K. and M.L. analyzed the results from mammalian cells; C.K., A.V.K., J.M.C., and J.G.Z. wrote the manuscript with input from all authors.

## Data and code availability

Screen data generated during this study will be deposited to GEO. Code using for bioinformatic analysis will be available on GitHub.

## Materials and methods

### Bacterial strains and plasmid constructs

*E. coli* K-12 substrain MG1655 was used for all CRISPRa tests unless specified. CD38 with highly expressed mRFP was used for CRISPRi experiments (**Supplementary Table S1**). Plasmid constructs were cloned using standard molecular biology methods (Fontana et al., 2020a; Kiattisewee et al., 2021). All PCR fragments were amplified with Phusion DNA Polymerase (Thermo-Fisher Scientific) for Infusion Cloning (Takara Bio). Plasmids were transformed into chemically competent NEB Turbo *E. coli* (New England Biolabs) cells, plated on LB-agar, and cultured in LB media supplied with the appropriate antibiotics used in the following concentrations: 100 μg/mL Carbenicillin, 25 μg/mL Chloramphenicol, 30 μg/mL Kanamycin. All plasmid constructs were confirmed by Sanger sequencing (GENEWIZ).

PAM-flexible dCas9 variants were cloned from existing dCas9 and dxCas9(3.7) plasmids (pCD442 and pCD564) (Fontana et al., 2020a). dxCas9-NG was cloned by replacing the C-terminus of dxCas9(3.7) with dCas9-NG mutations ordered as a gBlock (IDT). dCas9-NG was cloned by fusing the N-terminus of dCas9 and C-terminus of dxCas9-NG together. dSpG and dSpRY sequences were cloned into a bacterial codon optimized vector (Addgene #101199) and then subcloned into the pCD442 vector. Complete sequences are provided in the supplementary information.

Each dCas9 variant (dCas9, dxCas9, dCas9-NG, dxCas9-NG, dSpG, and dSpRY) was expressed from the endogenous Sp.pCas9 promoter in a p15A vector (**Supplementary Table S2**). MCP-SoxS (R93A, S101A) (abbreviated MCP-SoxS) was expressed from the BBa_J23107 promoter (http://parts.igem.org) in the same plasmid with dCas9. The single guide RNAs (sgRNA) or modified scaffold RNAs b2.1xMS2 (scRNAs) were expressed from the strong BBa_J23119 promoter, either in the same plasmid with the dCas9-carrying plasmid or in a separate ColE1 plasmid (Dong et al., 2018; Fontana et al., 2020a). All 20 bp scRNA/sgRNA target sequences are provided in **Supplementary Table S3**. The mRFP reporter was expressed from the weak J3-BBa_J23117 promoter on a pSC101** plasmid (**Supplementary Table S2**). For CRISPRi experiments, a construct expressing mRFP from the strong BBa_J23119 promoter (strain CD38, **Supplementary Table S1**) was integrated into the *E. coli* genome using a previously-described lambda red system (Dong et al., 2018). For endogenous promoter CRISPRa experiments, we used GFPmut2 reporters on pSC101** vectors as described previously (Zaslaver et al., 2006). Reporters were purchased from Horizon Discovery or constructed with the same methodology (Zaslaver et al., 2006).

### Plate reader experiments

Single colonies from LB plates were inoculated in 400 μL of EZ-RDM (Teknova) supplemented with the appropriate antibiotics and grown in 96-deep-well plates at 37 °C with shaking overnight 900 RPM on a Heidolph titramax 1000. 150 μL of the overnight culture were transferred into flat, clear-bottomed black 96-well plates (Corning) and the OD_600_ and fluorescence were measured in a Biotek Synergy HTX plate reader. Data were analyzed using the BioTek Gen5 2.07.17 software. For mRFP1 detection, the excitation wavelength was 540 nm and emission wavelength was 600 nm. For GFPmut2 detection, the excitation wavelength was 485 nm and emission wavelength was 528 nm.

### Flow cytometry

Single colonies from LB plates were inoculated in 400 μL EZ-RDM (Teknova) supplemented with appropriate antibiotics and grown in 96-deep-well plates at 37 °C, 900 RPM on a Heidolph titramax 1000. Cultures were grown overnight and then diluted in 1:100 in Dulbecco’s phosphate-buffered saline (PBS) and analyzed on a MACSQuant VYB flow cytometer with the MACSQuantify 2.8 software (Miltenyi Biotec). To select single cells, we used a previously-described gating procedure (Dong et al., 2018). A side scatter threshold trigger (SSC-H) was applied to select for single cells until 10000 events were collected. FlowJo 10.0.7 software was used to apply a narrow gate along the diagonal line on the SSC-H vs SSC-A plot to exclude the events where multiple cells were grouped together. Within the selected population, events that appeared on the edges of the FSC-A vs. SSC-A plot and the fluorescence histogram were excluded.

### Pooled PAM library construction and screening

To generate the pooled reporter library with different PAMs at the target site -81 bp from the TSS, we used pJF143.J3 (Supplementary Table S2) as a PCR template with oCK679_NNN (5’-CTCGTCTCCTCACTTT<u>NNN</u>ACGGAGCGTTCTGGACACAACG-3’) as a forward primer and oCK680 (5’-AAGTGAGGAGACGAGCGAACGC-3’) as a reverse primer. Amplified linear fragments were treated with DpnI to remove the parental vector and circularized with Infusion. oCK679_NNN oligos with NGH, NHG, NHH, NRN, and NYN were used to construct each corresponding PAM library. For each screened variant, the number of colonies picked was 2X the number of sequence variants. For example, we picked 24 colonies for NGH (12 possible sequences) and 64 colonies for NRN (32 possible sequences). Fold activation was calculated relative to a strain with pJF143.J3 and an off-target scRNA.

### CRISPRa at endogenous promoters

CRISPRa at endogenous promoters was performed with a three plasmid system — dCas9 plasmid, scRNA plasmid, and reporter plasmid. Four promoters tested here (yajGp, uxuRp, araEp, and ppiDp2) were evaluated previously with dxCas9(3.7) (Fontana et al., 2020a). These promoters were chosen based on the criteria: 1) genes should not be highly expressed, and 2) genes should be regulated by the sigma70 family (Fontana et al., 2020a). A second set of endogenous promoters (aroKp1, aroLp, pheLp, secBp, aroFp, aroHp1, aroHp2, talAp, and serCp) were selected from genes involved in aromatic amino acid biosynthesis pathways that meet the same criteria described above. scRNAs for each promoter were designed to target the optimal positions (-70 and -80 for the template strand and -81 and -91 for non-template strand relative to the TSS). One scRNA from each strand was chosen, based on which PAM was predicted to be accessible to the highest number of dCas9 variants. Accessibility was assessed based on the moderate performance threshold cutoff (see Supplemental Table S4 and Supplementary Methods).

### Bioinformatic analysis of targetable genes

Previously reported data for the activity of PAM-flexible dCas9 variants were used to predict the targetable genes for each dCas9 variant. To identify the compatible PAMs for each engineered dCas9 variant, we used data from Cas9 nuclease assays for each variant (Kim et al., 2020; Legut et al., 2020; Walton et al., 2020). Predicted compatible PAMs for each variant are provided in **Supplementary Table S4** (see Supplementary Methods for further details). We then examined *E. coli* and *P. putida* genome sequences for PAM availability. For *E. coli*, 1265 omoters with strong confidence in TSS were retrieved from RegulonDB (Santos-Zavaleta et al., 2019). For *P. putida*, 1104 experimentally-confirmed primary transcriptional units were used for the analysis (D’Arrigo et al., 2016). The PAMs at optimal target site positions were retrieved: -70 and -80 for the template strand and -81 and -91 for non-template strands relative to the TSS. The promoters with at least one compatible PAM out of four target sites were considered targetable. Further information can be found in **Figure S3**.

## Data analysis

Flow cytometry analysis was conducted on FlowJo 10.0.7 software or Python FlowCytometryTools on Jupyter Notebooks and then further processed with Microsoft Excel and Graphpad Prism. Data represents the average and standard deviation of at least three biologically-independent replicates, unless specified. Fold activation and fold repression were calculated by comparing sample fluorescence with a strain expressing off-target scRNA/sgRNA.

