## Supporting Information for "Expanding the scope of bacterial CRISPR activation with PAM-flexible dCas9 variants"

#: current affiliation: Beam Therapeutics
Cambridge, MA 02142
United States

+: These authors contributed equally
*: Corresponding authors

206-221-4902


206-543-1670

#### Table of Contents

**Supplementary Figures** 4

**Figure S1: Mutations contributing to PAM-flexibility of engineered dCas9 variants** 4

**Figure S2: CRISPRko and CRISPRa in mammalian systems with PAM-flexible dCas9 variants** 5

**Figure S3: PAM availability and distribution in E. coli and P. putida promoters** 6

**Figure S4: Basal expression of the mRFP reporter gene with different dCas9 variants and PAMs sequences** 7

**Figure S5: CRISPRi with PAM-flexible dCas9 variants** 8

**Figure S6: CRISPRa at endogenous promoters with PAM-flexible dCas9 variants** 9

**Supplementary Tables** 11

**Supplementary Table S1: E. coli strains.** 11

**Supplementary Table S2: Selected E. coli plasmids.** 11

**Supplementary Table S3: scRNA and sgRNA target sites.** 13

**Supplementary Table S4: Predicted compatible PAMs for each dCas9 variants** 14

**Supplementary Table S5: CRISPRa on endogenous E. coli promoters** 17

**Supplementary Methods** 18

**Evaluating PAM accessibility from CRISPR knockout and CRISPR activation screens in mammalian systems** 18

**PAM compatibility analysis** 18

**Endogenous promoter and scRNA selection strategy** 19

**DNA sequences** 20

**References** 30

#### Supplementary Figures

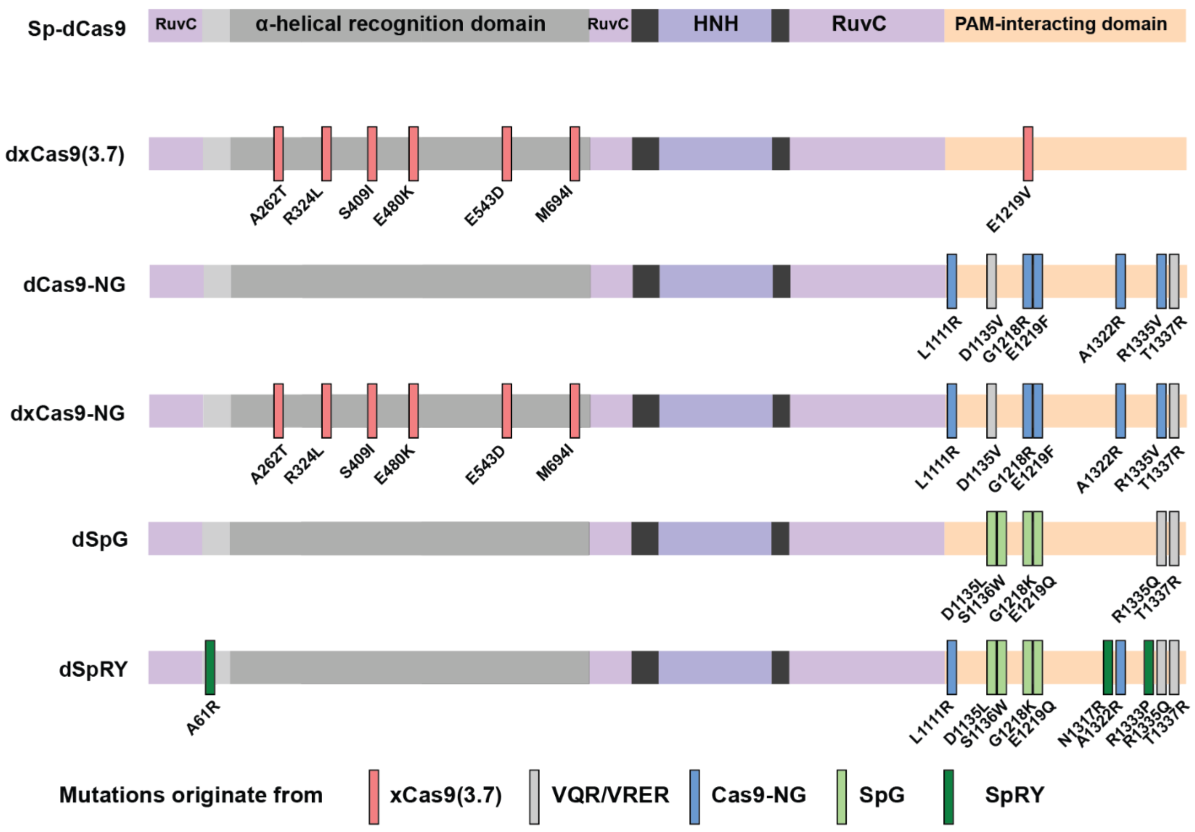


**Figure S1: Mutations contributing to PAM-flexibility of engineered dCas9 variants**

PAM-flexible dCas9 variants contain differing sets of mutations. All variants share the catalytically-inactivating D10A and H840A mutations of dCas9 (Qi et al., 2013). Mutations of dxCas9(3.7), from phage-assisted continuous evolution (PACE), were mostly at the α-helical recognition domain (Hu et al., 2018). dCas9-NG engineering was guided by the crystal structure with mutations made in the PAM-interacting domain (Nishimasu et al., 2018). dxCas9-NG has combined mutations from dxCas9(3.7) and dCas9-NG (Legut et al., 2020). dSpG and dSpRY were developed through structure-guided engineering with engineering trajectories monitored using a high-throughput PAM determination assay (Walton et al., 2021, 2020). SpG was engineered using mutations from the VRQR variant as a starting point (Kleinstiver et al., 2016, 2015), and SpRY contains mutations from Cas9-NG (Nishimasu et al., 2018). dSpRY is an evolved version of dSpG with a preference for NRN and some NYN PAMs (R is purine nucleotides, Y is pyrimidine nucleotides) (Walton et al., 2020). Mutation colors indicate the origin of each specific mutation. Complete sequences are provided in the DNA sequences section below.

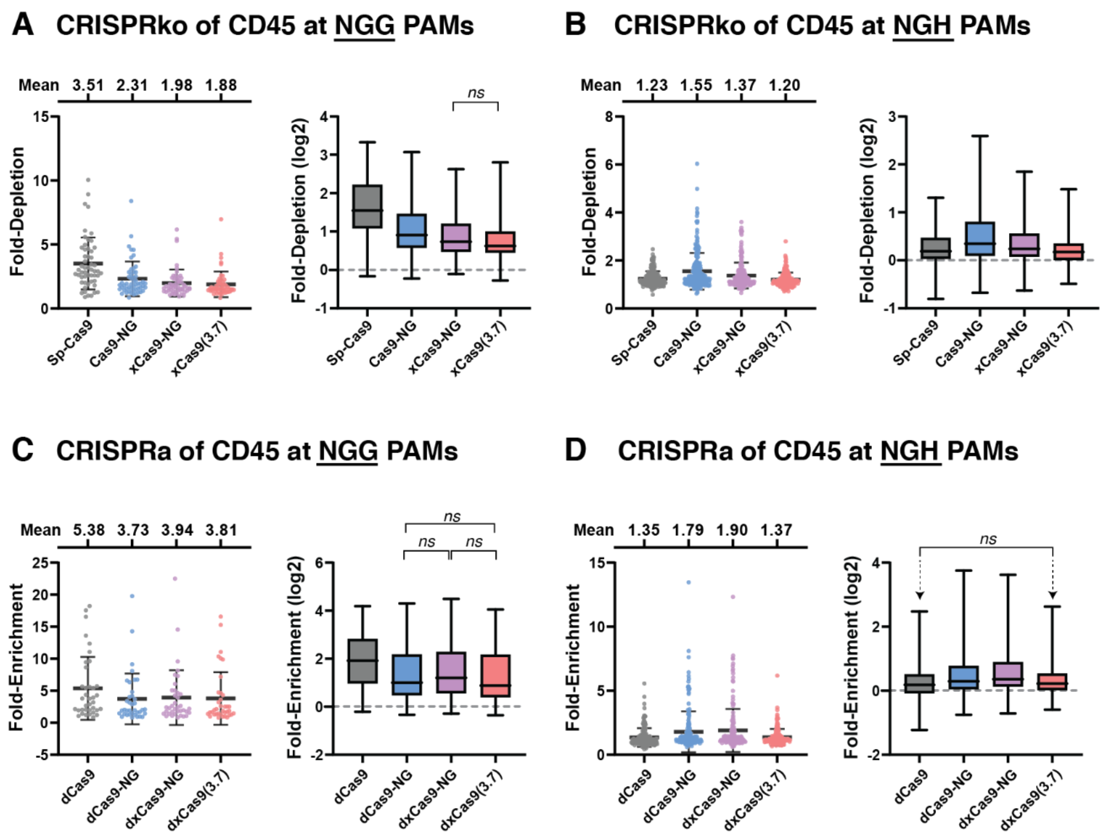


**Figure S2: CRISPRko and CRISPRa in mammalian systems with PAM-flexible dCas9 variants**

Cas9 variants were evaluated for CRISPRko (knockout) and CRISPRa (activation) at different PAMs. sgRNA-Cas9 effector complexes were targeted to the CD45 gene and changes in expression levels were evaluated by FACS-Seq (see Supplemental Methods). xCas9-NG and Cas9-NG exhibit comparable levels of PAM flexibility and both outperform xCas9(3.7). Data were plotted as fold-change scatter plots (left panel) or log2 fold-change box plots (right panel) for (A) CRISPRko at NGG PAMs, (B) CRISPRko at NGH PAMs, (C) CRISPRa at NGG PAMs, and (D) CRISPRa at NGH PAMs. In the scatter plots, bars and whiskers represent mean and standard deviation, respectively. In the box plots, two-tailed unpaired Welch's *t* test for each dCas9 pair were performed. Only non-significant comparisons (*ns*, $p > 0.05$) are indicated; all other differences (between proteins, within modalities) are significant.

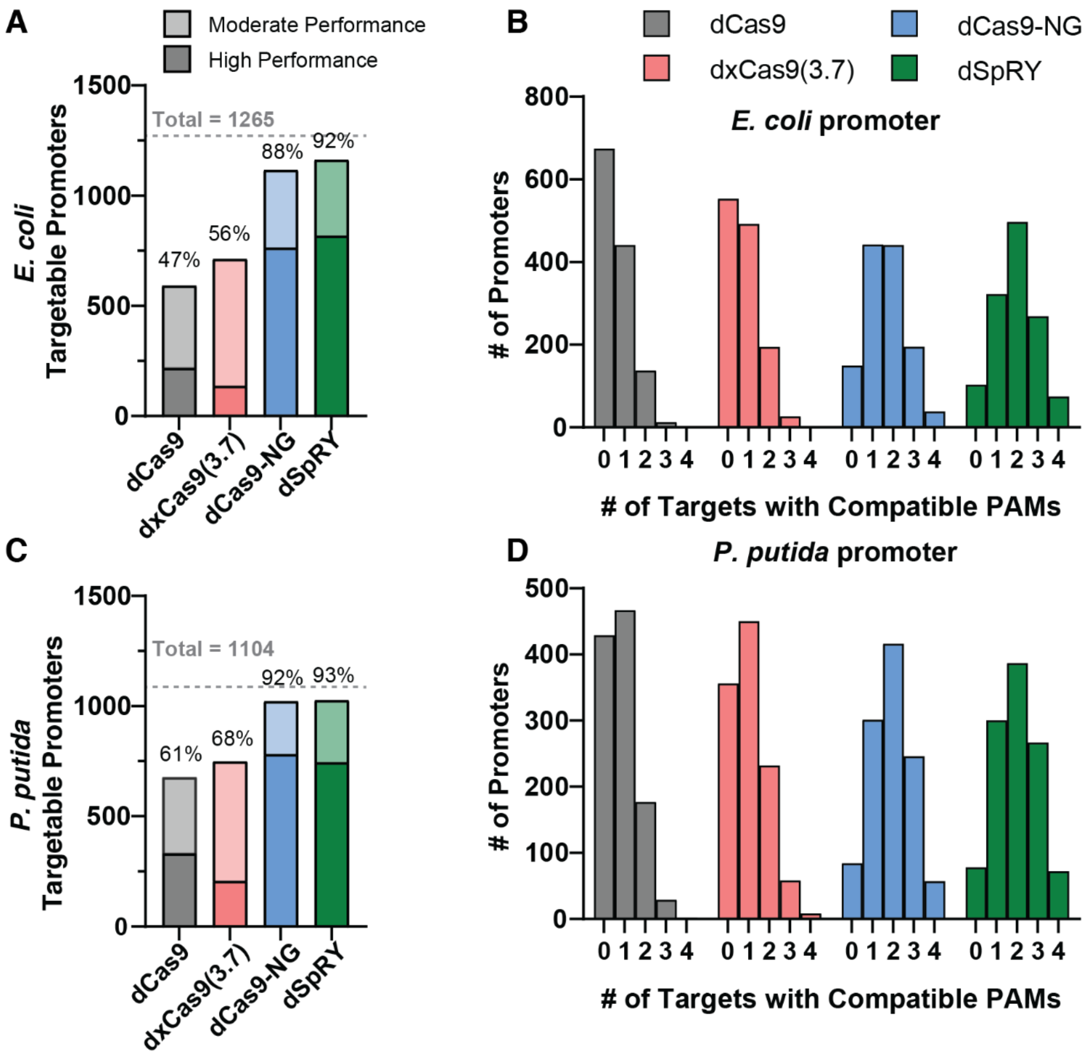


**Figure S3: PAM availability and distribution in *E. coli* and *P. putida* promoters**

Endogenous *E. coli* and *P. putida* promoters were analyzed for predicted targetable sites with different PAM-flexible dCas9 variants (see Methods section for further details). (A) Number of targetable *E. coli* promoters for different PAM-flexible dCas9 variants. 1265 *E. coli* promoters were analyzed for the presence of a compatible PAM at any of the four effective target site positions upstream of the TSS (see Methods). "High performance" PAMs are classified as PAMs with at least 50% efficiency relative to dCas9 at NGG PAMs. "Moderate performance" PAMs are classified as PAMs with 20-50% efficiency. (B) Distribution of promoters with 0 to 4 compatible PAMs with at least moderate performance in *E. coli*. (C) Number of targetable *P. putida* promoters for different PAM-flexible dCas9 variants. 1104 *P. putida* were analyzed with the same strategy. (D) Distribution of promoters with 0 to 4 compatible PAMs with at least moderate performance in *P. putida*. In both *E. coli* and *P. putida*, dxCas9-NG and dSpRY access more endogenous promoters than dCas9 and dxCas9(3.7). As *P. putida* has a higher GC content than *E. coli* and a correspondingly higher fraction of NGN PAMs, there is an almost identical number of targetable promoters for dCas9-NG and dSpRY.

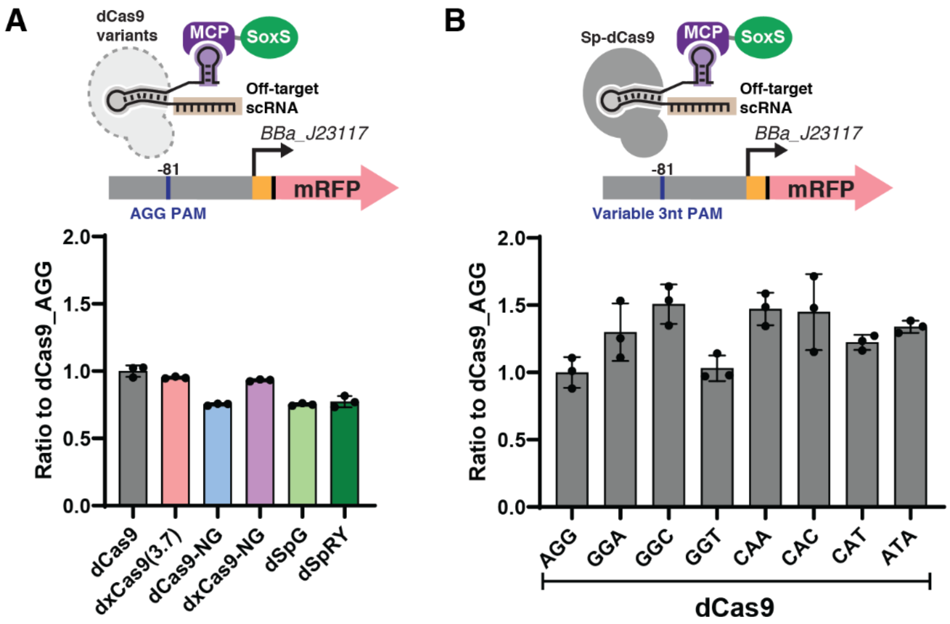


**Figure S4: Basal expression of the mRFP reporter gene with different dCas9 variants and PAMs sequences**

(A) Expression of different dCas9 variants produces less than 2-fold changes in basal reporter expression level. This experiment was performed with AGG PAM reporter and an off-target scRNA (hAAVS1). These off-target expression levels were used as a basal expression for fold-change calculation with each dCas9 variant in Figure 2. (B) Modified mRFP reporter genes with alternative upstream PAM sites produce less than 2-fold changes in basal reporter expression. This experiment was performed with Sp-dCas9 and an off-target scRNA (hAAVS1). Values in panel A and B represent the mean ± standard deviation calculated from n = 3.

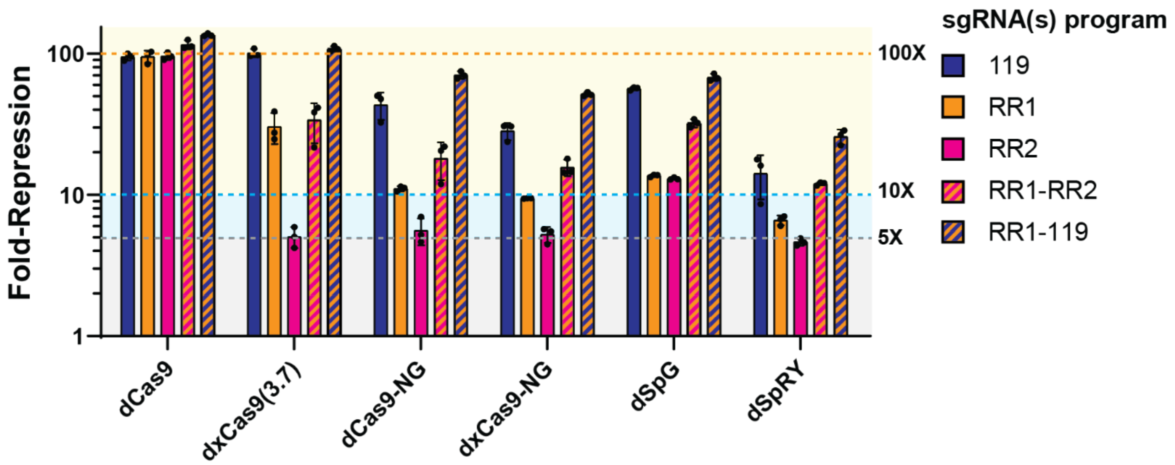


**Figure S5: CRISPRi with PAM-flexible dCas9 variants**

CRISPRi for PAM-flexible dCas9 variants was tested on an mRFP reporter gene with sgRNAs targeting the promoter (119) or coding sequence (RR1 and RR2) with one or two sgRNAs expressed, each targeting sites harboring NGG PAMs. dCas9 exhibited the highest repression efficiency (>90-fold) among all variants. When only a single sgRNA is expressed, the sgRNA targeting the promoter region (119) produces the largest repression effect with all dCas9 variants. The addition of the second sgRNA (RR1-RR2 or RR1-119) led to significant improvement in repression for all variants. Values represent the mean ± standard deviation calculated from n = 3.

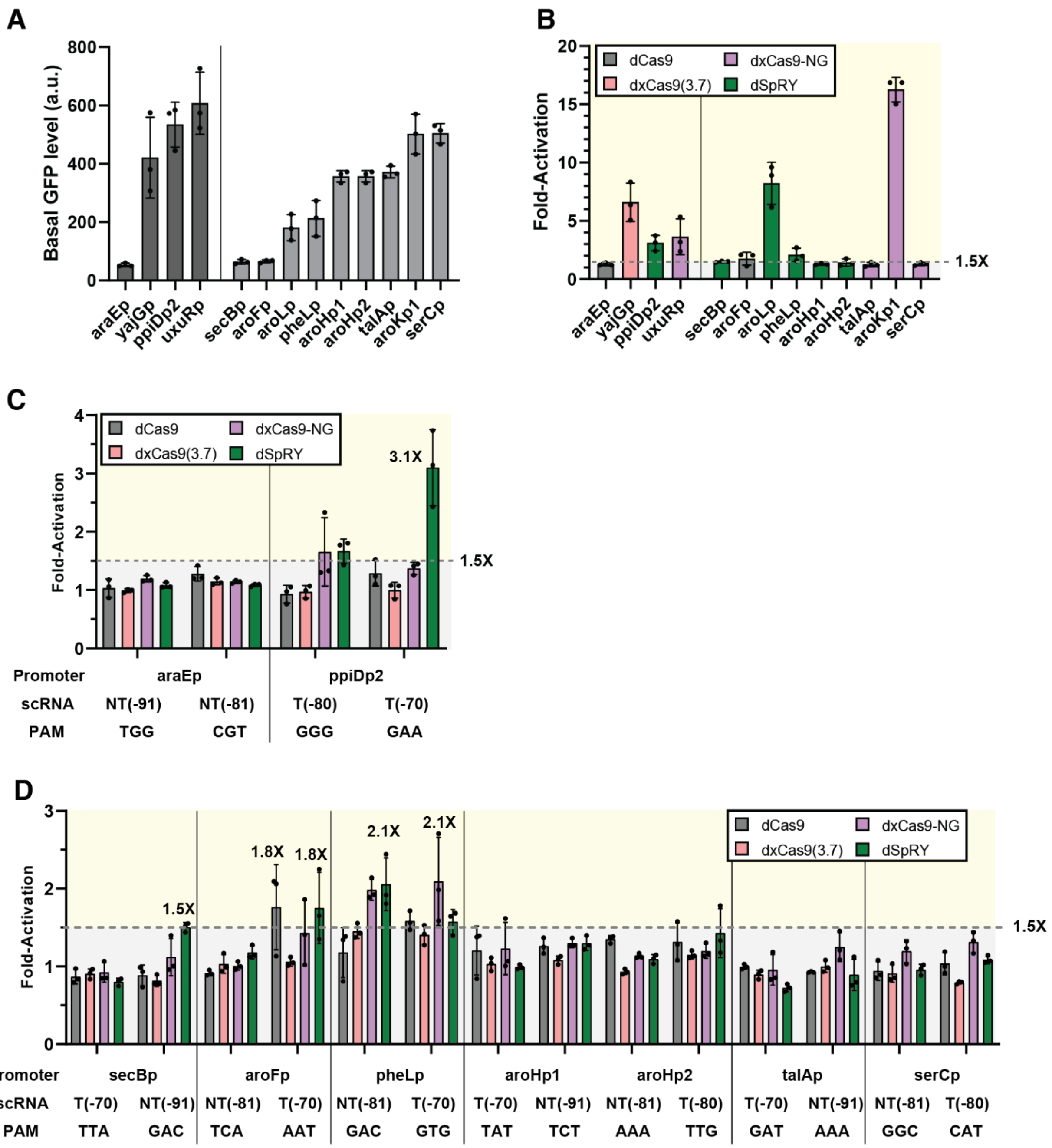


**Figure S6: CRISPRa at endogenous promoters with PAM-flexible dCas9 variants**

(A) Basal expression levels for endogenous promoters (left: promoters previously tested with dxCas9(3.7) (Fontana et al., 2020); right: new promoters examined in this work). Basal expression levels were measured in a strain expressing the parent dCas9 and an off-target scRNA (hAAVS1). (B) Maximum fold-activation for each endogenous promoter. Bar color indicates the dCas9 variant with the highest fold-activation for the corresponding endogenous promoter. 8 out of 13 endogenous promoters can be activated (>1.5-fold) with PAM-flexible dCas9 variants. (C) CRISPRa at endogenous promoters previously tested with dxCas9(3.7) (araEp and ppiDp2) using NGG or non-NGG PAMs. See main text Figure 4B for the other two promoters from this set (yajGp and uxuRp). (D) CRISPRa at endogenous promoters involved in aromatic amino acid biosynthesis using non-NGG PAMs (secBp, aroFp, pheLp, aroHp1, aroHp2, talAp, and serCp). See main text Figure 4C for the other two promoters from the aromatic amino acid biosynthesis set (aroLp and aroKp1). Data were collected by flow cytometry and fold-activation was calculated compared to the strain expressing the corresponding dCas9 variant and an off-target hAAVS1 scRNA. Values in panels A-D represent the mean ± standard deviation calculated from n = 3.

#### Supplementary Tables

**Supplementary Table S1: *E. coli* strains.**

| Strain | Description | Genotype |
| --- | --- | --- |
| MG1655 | Wildtype *E. coli strain* | F- λ- ilvG- rfb-50 rph-1 |
| CD38 | Integrated BBa_J23119-mRFP used for CRISPRi | MG1655 *nsfA*::BBa_J23119-mRFP |

**Supplementary Table S2: Selected *E. coli* plasmids.**

| Plasmid | Marker | Origin | Promoter | Gene | Terminator | Reference |
| --- | --- | --- | --- | --- | --- | --- |
| pCD442 | CmR | p15A | 1) Sp.pCas9<br>2) BBa_J23107 | 1) dCas9<br>2) MCP-SoxS | 1) BBa_B0015<br>2) BBa_B1002 | (Fontana et al., 2020) |
| pCD564 | CmR | p15A | 1) Sp.pCas9<br>2) BBa_J23107 | 1) dxCas9(3.7)<br>2) MCP-SoxS | 1) BBa_B0015<br>2) BBa_B1002 | (Fontana et al., 2020) |
| pCK668 | CmR | p15A | 1) Sp.pCas9<br>2) BBa_J23107 | 1) dCas9-NG<br>2) MCP-SoxS | 1) BBa_B0015<br>2) BBa_B1002 | This work |
| pCK669 | CmR | p15A | 1) Sp.pCas9<br>2) BBa_J23107 | 1) dxCas9-NG<br>2) MCP-SoxS | 1) BBa_B0015<br>2) BBa_B1002 | This work |
| pCK340 | CmR | p15A | 1) Sp.pCas9<br>2) BBa_J23107 | 1) dSpG<br>2) MCP-SoxS | 1) BBa_B0015<br>2) BBa_B1002 | This work |
| pCK341 | CmR | p15A | 1) Sp.pCas9<br>2) BBa_J23107 | 1) dSpRY<br>2) MCP-SoxS | 1) BBa_B0015<br>2) BBa_B1002 | This work |
| pCK085.scRNA | CmR | p15A | 1) Sp.pCas9<br>2) BBa_J23107<br>3) BBa_J23119 | 1) dCas9<br>2) MCP-SoxS<br>3) scRNA | 1) BBa_B0015<br>2) BBa_B1002<br>3) TrrnB | (Tickman et al., 2021) |
| pCK281.scRNA | CmR | p15A | 1) Sp.pCas9<br>2) BBa_J23107<br>3) BBa_J23119 | 1) dxCas9(3.7)<br>2) MCP-SoxS<br>3) scRNA | 1) BBa_B0015<br>2) BBa_B1002<br>3) TrrnB | This work |
| pCK670.scRNA | CmR | p15A | 1) Sp.pCas9<br>2) BBa_J23107<br>3) BBa_J23119 | 1) dCas9-NG<br>2) MCP-SoxS<br>3) scRNA | 1) BBa_B0015<br>2) BBa_B1002<br>3) TrrnB | This work |
| pCK671.scRNA | CmR | p15A | 1) Sp.pCas9<br>2) BBa_J23107<br>3) BBa_J23119 | 1) dxCas9-NG<br>2) MCP-SoxS<br>3) scRNA | 1) BBa_B0015<br>2) BBa_B1002<br>3) TrrnB | This work |
| pCK363.scRNA | CmR | p15A | 1) Sp.pCas9<br>2) BBa_J23107<br>3) BBa_J23119 | 1) dSpG<br>2) MCP-SoxS<br>3) scRNA | 1) BBa_B0015<br>2) BBa_B1002<br>3) TrrnB | This work |
| pCK364.scRNA | CmR | p15A | 1) Sp.pCas9<br>2) BBa_J23107<br>3) BBa_J23119 | 1) dSpRY<br>2) MCP-SoxS<br>3) scRNA | 1) BBa_B0015<br>2) BBa_B1002<br>3) TrrnB | This work |

| | | | | | | |
|---|---|---|---|---|---|---|
| pJF143. J3 | AmpR | pSC101** | J3-BBa_J23117 | mRFP | BBa_B0015 | (Fontana et al., 2020) |
| pCK284. NNN | AmpR | pSC101** | J3(NNN-PAM)-BBa_J23117 | mRFP | BBa_B0015 | This work |
| pCD443. scRNA | AmpR | ColE1 | BBa_J23119 | scRNA | TrrnB | This work |
| pCK411. scRNA/ sgRNA | AmpR | ColE1 | BBa_J23119 | scRNA/sgRNA | BBa_K2680405 | This work |
| pCK590. XXX | KmR | pSC101 | Endogenous (strand +) | GFPmut2 | | Same as (Zaslaver et al., 2006) |
| pCK591. XXX | KmR | pSC101 | Endogenous (strand –) | GFPmut2 | | Same as (Zaslaver et al., 2006) |

**Supplementary Table S3: scRNA and sgRNA target sites.**

| sc/sgRNA | DNA sequence | Target strand[a] | Distance to TSS[b] | PAM |
|---|---|---|---|---|
| J306 | TTGTGTCCAGAACGCTCCGT | NT | -81 | AGG |
| hAAVS1 | GGGGCCACTAGGGACAGGAT | Off-target | NA | – |
| RR1 | AACTTTCAGTTTAGCGGTCT | NT | 151[c] | GGG |
| RR2 | TGGAACCGTACTGGAACTGC | NT | 215[c] | GGG |
| 119 | AATTCAGATCTATTATACCT | NT | -15 | AGG |
| yajGp_Y1 | TTGACGAAATAATCGCCCCT | NT | -81 | GGT |
| yajGp_Y2 | CATCAGTGTTTCTTTTACCA | T | -79 | GGG |
| uxuRp_U1 | TGATTGACCAGTAAGTCTGT | NT | -81 | AGG |
| uxuRp_U2 | GATTACCCTACAGACTTACT | T | -70 | GGT |
| ppiDp2_D1 | ACTAAGCGTTGTCCCCAGTG | T | -80 | GGG |
| ppiDp2_D2 | GTCCCCAGTGGGGATGTGAC | T | -70 | GAA |
| araEp_E1 | TGCGACATGTCGTTATGTGA | NT | -91 | TGG |
| araEp_E2 | ATTAAATTGCTGCGACATGT | NT | -81 | CGT |
| pheLp_X06 | GCGATACACTCAATATAAAG | NT | -81 | GAC |
| pheLp_X07 | AGAGTAGTCCTTTATATTGA | T | -70 | GTG |
| secBp_X07 | CACCACGGTTCCCCAGATTT | T | -70 | TTA |
| secBp_X08 | AATCTGGGGAACCGTGGTGC | NT | -91 | GAC |
| serCp_X06 | ACCGTTGAGGGCAAAAATGT | NT | -81 | GGC |
| serCp_X09 | CTTTTGTGTGATGCAAGCCA | T | -80 | CAT |
| talAp_X07 | GGTAATAATCCTATAACACT | T | -70 | GAT |
| talAp_X08 | GTGTTATAGGATTATTACCA | NT | -91 | AAA |
| aroFp_X06 | CAGGCAATTTAGTCGCGCTT | NT | -81 | TCA |
| aroFp_X07 | AAGGGTTGAAAGCGCGACTA | T | -70 | AAT |
| aroLp_X07 | TGGTGGCTGGAAGTGCAACG | T | -70 | TAG |
| aroLp_X08 | GTTGCACTTCCAGCCACCAC | NT | -91 | TTC |
| aroHp1_X07 | ATTGCCACCAAGATCCTCGA | T | -70 | TAT |
| aroHp1_X08 | CGAGGATCTTGGTGGCAATC | NT | -91 | TCT |
| aroHp2_X06 | GTGGTTAGCATGATAACAAA | NT | -81 | AAA |
| aroHp2_X09 | TAGTGCATTAGCTTATTTTT | T | -80 | TTG |
| aroKp1_X07 | GAGTAAACAGCCGTAAAAGC | T | -70 | GGT |
| aroKp1_X08 | CTTTTACGGCTGTTTACTCA | NT | -91 | CTG |

[a] *Template strand (T) or non-template strand (NT).*
[b] *Distance from the 3' end of the guide site (PAM proximal) to the TSS. For synthetic promoters (BBa_J23117 or BBa_J23119, http://parts.igem.org), the TSS is immediately downstream.*
[c] *For RR1 and RR2 sgRNAs, the positive number refers to the distance downstream of the TSS.*

**Supplementary Table S4: Predicted compatible PAMs for each dCas9 variants**

| PAM | Sp-Cas9 | xCas9(3.7) | Cas9-NG | SpG | SpRY |
|---|---|---|---|---|---|
| AAA | 0.18% | 2.75% | 19.51% | N.A. | 27.97% |
| AAC | 0.06% | 1.52% | 17.62% | N.A. | 69.73% |
| AAG | 46.32% | 19.04% | 43.10% | N.A. | 28.35% |
| AAT | 0.25% | 3.80% | 28.04% | N.A. | 6.58% |
| ACA | 0.61% | 0.23% | 5.03% | N.A. | 19.84% |
| ACC | -0.50% | -0.15% | 2.18% | N.A. | N.A. |
| ACG | 17.85% | 7.43% | 12.10% | N.A. | N.A. |
| ACT | -0.45% | 0.47% | 7.07% | N.A. | N.A. |
| AGA | 27.95% | 29.33% | 60.81% | 65.89% | 58.50% |
| AGC | 7.82% | 16.70% | 52.14% | 75.97% | 56.19% |
| AGG | 99.80% | 62.37% | 72.89% | 38.75% | 21.46% |
| AGT | 1.82% | 23.39% | 68.90% | 78.02% | 59.55% |
| ATA | 0.78% | 0.21% | 12.29% | N.A. | 43.63% |
| ATC | -0.56% | -0.34% | 5.07% | N.A. | 3.20% |
| ATG | 22.04% | 5.45% | 28.21% | N.A. | N.A. |
| ATT | -0.13% | 0.55% | 10.60% | N.A. | N.A. |
| CAA | -0.01% | 6.21% | 27.29% | N.A. | 53.17% |
| CAC | -0.53% | 2.69% | 26.45% | N.A. | 19.04% |
| CAG | 44.62% | 25.71% | 47.15% | N.A. | 18.17% |
| CAT | -0.63% | 6.25% | 32.92% | N.A. | 22.44% |
| CCA | -0.13% | -0.21% | 2.06% | N.A. | N.A. |
| CCC | -0.65% | -0.31% | 0.89% | N.A. | 1.01% |
| CCG | 11.47% | 3.99% | 6.78% | N.A. | 9.06% |
| CCT | -0.22% | -0.37% | 3.09% | N.A. | N.A. |

| | | | | | |
|---|---|---|---|---|---|
| CGA | 30.35% | 36.77% | 59.10% | 78.85% | N.A. |
| CGC | 10.29% | 24.23% | 52.31% | 70.58% | N.A. |
| CGG | 102.08% | 68.15% | 72.25% | 48.14% | 15.80% |
| CGT | 2.47% | 32.69% | 69.50% | 49.23% | 23.77% |
| CTA | -0.23% | 0.64% | 9.53% | N.A. | 20.11% |
| CTC | -0.52% | 0.19% | 4.65% | N.A. | 25.22% |
| CTG | 17.31% | 3.94% | 23.82% | N.A. | 19.44% |
| CTT | -0.44% | -0.06% | 9.93% | N.A. | 2.30% |
| GAA | 1.02% | 11.96% | 37.31% | N.A. | 100.83% |
| GAC | -0.13% | 4.71% | 31.57% | N.A. | 86.73% |
| GAG | 53.38% | 35.04% | 57.91% | N.A. | 55.14% |
| GAT | 0.05% | 10.29% | 42.64% | N.A. | 35.73% |
| GCA | -0.05% | 0.76% | 7.21% | N.A. | 55.08% |
| GCC | -0.26% | 0.28% | 4.37% | N.A. | 21.67% |
| GCG | 13.01% | 1.67% | 11.22% | N.A. | N.A. |
| GCT | -0.58% | 0.22% | 10.36% | N.A. | 43.08% |
| GGA | 30.31% | 37.93% | 60.78% | 62.42% | 33.41% |
| GGC | 12.24% | 22.95% | 53.55% | 73.33% | 41.75% |
| GGG | 101.50% | 70.07% | 76.36% | 67.74% | 54.37% |
| GGT | 7.16% | 33.76% | 69.40% | 76.14% | 65.25% |
| GTA | 0.05% | 1.79% | 20.16% | N.A. | 37.98% |
| GTC | -0.34% | 0.56% | 11.00% | N.A. | N.A. |
| GTG | 23.36% | 6.76% | 37.60% | N.A. | 50.91% |
| GTT | -0.15% | 0.29% | 16.20% | N.A. | 4.66% |
| TAA | 0.62% | 1.36% | 16.05% | N.A. | 69.86% |
| TAC | -0.62% | 0.15% | 11.41% | N.A. | 72.28% |

| | | | | | |
|---|---|---|---|---|---|
| **TAG** | 36.56% | 11.45% | 34.32% | N.A. | 30.23% |
| **TAT** | -0.22% | 1.52% | 19.26% | N.A. | 44.40% |
| **TCA** | -0.36% | 0.01% | 1.27% | N.A. | N.A. |
| **TCC** | -0.81% | -0.28% | 0.72% | N.A. | N.A. |
| **TCG** | 10.87% | 3.10% | 5.79% | N.A. | N.A. |
| **TCT** | -0.15% | -0.37% | 2.47% | N.A. | 7.00% |
| **TGA** | 28.62% | 29.68% | 61.71% | 73.33% | 45.34% |
| **TGC** | 7.91% | 15.40% | 53.76% | 88.36% | 65.05% |
| **TGG** | 96.62% | 62.22% | 72.08% | 95.59% | 69.12% |
| **TGT** | 2.32% | 24.78% | 70.19% | 103.69% | N.A. |
| **TTA** | 0.06% | 0.53% | 3.71% | N.A. | N.A. |
| **TTC** | -1.16% | -0.09% | 0.44% | N.A. | N.A. |
| **TTG** | 9.19% | 2.01% | 11.05% | N.A. | 0.14% |
| **TTT** | -0.46% | -0.14% | 1.55% | N.A. | 11.70% |

The predicted PAM compatibility of each PAM-Cas9 pair was reported as indel frequency (in % units) relative to the benchmark efficiency of Sp-Cas9 at NGG PAMs from each experiment (see Supplementary Methods) (Kim et al., 2020; Walton et al., 2020). Each data point was color-coded based on predicted efficiency: Green (high efficiency, >50% of benchmark), blue (moderate efficiency, 20%–50%), and gray (low efficiency, less than 20%). NGG PAMs are highlighted in yellow. N.A. means data is not available. Some of the reported efficiency values are small negative numbers, likely due to negligible editing frequencies indistinguishable from the background. Comparable data for indel frequencies with xCas9-NG are not available; for this work we assumed Cas9-NG and xCas9-NG were similar based on their comparable performance in nuclease assays (Figure S2).

**Supplementary Table S5: CRISPRa on endogenous *E. coli* promoters**

| Promoter | Max. FA | dCas9 variant | Position | PAM |
|---|---|---|---|---|
| **yajGp** | **6.6**<br>6.2<br>5.1 | **dxCas9(3.7)**<br>dxCas9-NG<br>dSpRY | NT(-81) | GGT |
| **uxuRp** | **3.6**<br>2.6 | **dxCas9-NG**<br>dSpRY | T(-70) | GGT |
| araEp | 1.3 | dxCas9-NG | NT(-81) | CGT |
| **ppiDp2** | **3.1** | **dSpRY** | T(-70) | GAA |
| **aroKp1** | **16.3**<br>12.8 | **dxCas9-NG**<br>dxCas9(3.7) | T(-70) | GGT |
| **aroLp** | **8.2**<br>5.5 | **dSpRY**<br>dCas9 | T(-70) | TAG |
| **aroF** | **1.8**<br>**1.8** | **dCas9**<br>**dSpRY** | T(-70) | AAT |
| aroHp1 | 1.3 | dxCas9-NG | NT(-91) | TCT |
| aroHp2 | 1.4 | dSpRY | NT(-81) | AAA |
| **pheLp** | **2.1**<br>**2.1** | **dxCas9-NG**<br>**dSpRY** | T(-70)<br>NT(-81) | GTG<br>GAC |
| **secBp** | **1.5** | **dSpRY** | NT(-91) | GAC |
| serCp | 1.3 | dxCas9-NG | T(-80) | CAT |
| talAp | 1.2 | dxCas9-NG | NT(-91) | AAA |

The top two conditions were shown for each promoter, if >1.5-fold. The best conditions for each promoter is shown in bold. yajGp has three similar activation levels. Promoters with <1.5-fold activation were shaded in gray.

### Supplementary Methods

#### Evaluating PAM accessibility from CRISPR knockout and CRISPR activation screens in mammalian systems

To predict the PAM accessibility for different Cas9 variants, we used data from CRISPR knockout (CRISPRko) and CRISPR activation (CRISPRa) screens in mammalian cells. Previously, screening data has been reported for three Cas9 variants (Sp-Cas9, xCas9(3.7), and Cas9-NG) (Legut et al., 2020). We report here a corresponding screen performed using the same approach with four Cas9 variants: Sp-Cas9, xCas9(3.7), Cas9-NG, and xCas9-NG. Lentiviral sgRNA libraries were constructed to target the CD45 gene, spanning the 3 kb region surrounding the TSS and constitutive CDS exons, using all possible 20-mer sequences upstream of an NG PAM sequence, as described before (Legut et al., 2020). In addition to sgRNAs, each vector also contained a Cas9 variant (nuclease for CRISPRko and catalytically dead Cas9 fusion with VPR transcriptional activators for CRISPRa) and a short barcode downstream of the sgRNA to identify the Cas9 variant in the same Illumina read as the sgRNA. sgRNA library cloning, lentivirus production and cell transduction were done separately for each Cas9 variant. The transduced and selected cells were only pooled together prior to FACS sorting based on the CD45 protein expression. CRISPRko was performed in K562 cells and CRISPRa was performed in A375 cells; both cell lines were obtained from ATCC. The presort samples and 10% top/bottom bins were analyzed by high-throughput Illumina sequencing. To calculate fold-depletion for CRISPRko, the relative frequency (normalized to sequencing depth and median frequency of non-targeting sgRNAs for each Cas9 variant) of each sgRNA from the bottom 10% bin (lowest expression level) was divided by its corresponding frequency in the top 10% bin. To calculate fold-enrichment for CRISPRa, the relative frequency of each sgRNA from the top 10% bin (highest expression) was divided by its corresponding frequency in the bottom 10% bin. For CRISPRko, only sgRNAs targeting within the CDS were included in the analysis; for CRISPRa, only sgRNAs targeting within the core promoter region (as determined previously (Legut et al., 2020), chr1:198638250-198639226) were included. The data were then filtered for designated PAMs (NGG or NGH) to visualize the data as shown in Figure S2.

#### PAM compatibility analysis

For each dCas9 variant, the ability to target each of the 64 possible 3-nucleotide PAMs was predicted based on previously reported CRISPR nuclease (CRISPRko) data in mammalian systems. (Kim et al., 2020; Walton et al., 2020). Genome editing efficiencies, measured as indel frequencies, have been reported for Sp-Cas9, xCas9(3.7), and Cas9-NG (Kim et al., 2020) and for Sp-Cas9, SpG, and SpRY (Walton et al., 2020). For each Cas9 dataset, sgRNAs with the same 3-nt PAM were grouped together. We then calculated an average indel frequency for all sgRNAs with the same PAM. We assessed PAM compatibility by comparison to the benchmark indel frequency of Sp-Cas9 at NGG PAMs (Table S4). “High performance” PAMs were those with >50% indel frequency relative to the benchmark. “Moderate performance” PAMs were those with 20-50% efficiency relative to the benchmark. PAMs that were not tested in the previously-reported nuclease assays (Walton et al., 2020) were assumed to be incompatible.

**Endogenous promoter and scRNA selection strategy**

Gene candidates were selected from metabolic pathways related to aromatic amino acid biosynthesis, including the relevant genes in central metabolic pathways. We selected only promoters that are regulated by sigma70 and are available in the previously characterized *E. coli* endogenous promoter library (obtained from Horizon Discovery) (Zaslaver et al., 2006). To identify potentially activatable promoters with moderately-weak basal expression levels, we compared basal expression to yajGp, which was the strongest endogenous promoter that could be activated in previous work (Fontana et al., 2020). We eliminated any promoters with >10-fold higher expression levels than yajGp, yielding 9 promoters: secBp, aroFp, aroLp, pheLp, aroHp1, aroHp2, talAp, aroKp1, and serCp. Some of these promoters regulate multi-gene operons. Complete sequences of the endogenous promoters used in this study are available in the DNA sequences section below.

Out of selected genes, aroH contains two putative promoters — aroHp1/aroHp2. In Figure S6A, the basal expression levels for aroHp1 and aroHp2 were assumed to be the same value. Expression levels produced by each individual promoter cannot be determined from the fluorescent reporter used in this study because both promoters are upstream of the same fluorescent reporter gene.

In the secBp promoter constructed for the *E. coli* promoter library (Zaslaver et al., 2006), only the sequence to -7 bases from the TSS was included. To include enough sequence for scRNA target sites, we constructed a promoter extending to -137 bases from the TSS.

For each promoter, four scRNAs were identified according to the previously described target site preference. X06 and X08 represent the scRNAs targeting the non-template strand at -81 and -91 positions (Table S3). X07 and X09 represent the scRNAs targeting the template strand at -70 and -80 positions. One target site from the template strand and another from the non-template strand were selected for further analysis based on which PAM was predicted to be accessible to the highest number of dCas9 variants. Accessibility was assessed based on the moderate performance threshold cutoff (Supplemental Table S4).

### DNA sequences

Reporter

>J3(**PAM**)-BBa_J23117-RBS-mRFP

AGCATTTGCGATCATTCACGCAGCGCTTATTCAGTTGCTCACTGCGATGTCATAATCATCGCTACGAGCTGTGAAAGAT
GCATAAAGCTCGTACGACGCGTTCGCTCGTCTCCTCACTTCT**NNN**ACGGAGCGTTCTGGACACAACGTCGTCTTGAAGT
TGCGATTATAGATTGACAGCTAGCTCAGTCCTAGGGATTGTGCTAGCGAATTCATTAAAGAGGAGAAAGGTACC**ATG**GC
GAGTAGCGAAGACGTTATCAAAGAGTTCATGCGTTTCAAAGTTCGTATGGAAGGTTCCGTTAACGGTCACGAGTTCGAA
ATCGAAGGTGAAGGTGAAGGTCGTCCGTACGAAGGTACCCAGACCGCTAAACTGAAAGTTACCAAAGGTGGTCCGCTGC
CGTTCGCTTGGGACATCCTGTCCCCGCAGTTCCAGTACGGTTCCAAAGCTTACGTTAAACACCCGGCTGACATCCCGGA
CTACCTGAAACTGTCCTTCCCGGAAGGTTTCAAATGGGAACGTGTTATGAACTTCGAAGACGGTGGTGTTGTTACCGTT
ACCCAGGACTCCTCCCTGCAAGACGGTGAGTTCATCTACAAAGTTAAACTGCGTGGTACCAACTTCCCGTCCGACGGTC
CGGTTATGCAGAAAAAAACCATGGGTTGGGAAGCTTCCACCGAACGTATGTACCCGGAAGACGGTGCTCTGAAAGGTGA
AATCAAAATGCGTCTGAAACTGAAAGACGGTGGTCACTACGACGCTGAAGTTAAAACCACCTACATGGCTAAAAAACCG
GTTCAGCTGCCGGGTGCTTACAAAACCGACATCAAACTGGACATCACCTCCCACAACGAAGACTACACCATCGTTGAAC
AGTACGAACGTGCTGAAGGTCGTCACTCCACCGGTGCTTAA

PAM location is bolded/underlined above

>BBa_J23119-RBS-mRFP

TTGACAGCTAGCTCAGTCCTAGGTATAATAGATCTGAATTCATTAAAGAGGAGAAAGGTACC**ATG**GCGAGTAGCGAAGA
CGTTATCAAAGAGTTCATGCGTTTCAAAGTTCGTATGGAAGGTTCCGTTAACGGTCACGAGTTCGAAATCGAAGGTGAA
GGTGAAGGTCGTCCGTACGAAGGTACCCAGACCGCTAAACTGAAAGTTACCAAAGGTGGTCCGCTGCCGTTCGCTTGGG
ACATCCTGTCCCCGCAGTTCCAGTACGGTTCCAAAGCTTACGTTAAACACCCGGCTGACATCCCGGACTACCTGAAACT
GTCCTTCCCGGAAGGTTTCAAATGGGAACGTGTTATGAACTTCGAAGACGGTGGTGTTGTTACCGTTACCCAGGACTCC
TCCCTGCAAGACGGTGAGTTCATCTACAAAGTTAAACTGCGTGGTACCAACTTCCCGTCCGACGGTCCGGTTATGCAGA
AAAAAACCATGGGTTGGGAAGCTTCCACCGAACGTATGTACCCGGAAGACGGTGCTCTGAAAGGTGAAATCAAAATGCG
TCTGAAACTGAAAGACGGTGGTCACTACGACGCTGAAGTTAAAACCACCTACATGGCTAAAAAACCGGTTCAGCTGCCG
GGTGCTTACAAAACCGACATCAAACTGGACATCACCTCCCACAACGAAGACTACACCATCGTTGAACAGTACGAACGTG
CTGAAGGTCGTCACTCCACCGGTGCTTAA

119, RR1, and RR2 are underlined above

>UP_secBp-UTR-RBS-GFPmut2

GATGGCAACGCCGCCAAGCGTGAAGAGATGATCAAACGCAGCGGTCGCACCACGGTTCCCCAGATTTTTATTGACGCAC
AGCACATTGGCGGCTGTGATGACTTGTATGCATTGGATGCACGTGGTGGACTGGATCCCCTGCTGAAATAACGTGTGAA
CGTTGGCATTACATTGCGCAGTATTTAAGGACAACACTTAAGGGTTTTCTACACATGTCAGAACAAAACAACACTGAAA
TGACTTTCCAGATCCAACGTATTTATACCAAGGATATCTCTTTCGAAGCGCCGCTCGAGAGATCCTCTAGATTTAAGAA
GGAGATATACAT**ATG**AGTAAAGGAGAAGAACTTTTCACTGGAGTTGTCCCAATTCTTGTTGAATTAGATGGTGATGTTA
ATGGGCACAAATTTTCTGTCAGTGGAGAGGGTGAAGGTGATGCAACATACGGAAAACTTACCCTTAAATTTATTTGCAC
TACTGGAAAACTACCTGTTCCATGGCCAACACTTGTCACTACTTTCGCGTATGGTCTTCAATGCTTTGCGAGATACCCA
GATCATATGAAACAGCATGACTTTTTCAAGAGTGCCATGCCCGAAGGTTATGTACAGGAAAGAACTATATTTTTCAAAG
ATGACGGGAACTACAAGACACGTGCTGAAGTCAAGTTTGAAGGTGATACCCTTGTTAATAGAATCGAGTTAAAAGGTAT
TGATTTTAAAGAAGATGGAAACATTCTTGGACACAAATTGGAATACAACTATAACTCACACAATGTATACATCATGGCA
GACAAACAAAAGAATGGAATCAAAGTTAACTTCAAAATTAGACACAACATTGAAGATGGAAGCGTTCAACTAGCAGACC
ATTATCAACAAAATACTCCAATTGGCGATGGCCCTGTCCTTTTACCAGACAACCATTACCTGTCCACACAATCTGCCCT
TTCGAAAGATCCCAACGAAAAGAGAGACCACATGGTCCTTCTTGAGTTTGTAACAGCTGCTGGGATTACCCATGGTATG
GATGAATTGTACAAATAA

Cas9 variants (bolded/underlined mutations)

>*Sp*.pCas9-dCas9-dblTerm

TTACGAAATCATCCTGTGGAGCTTAGTAGGTTTAGCAAGATGGCAGCGCCTAAATGTAGAATGATAAAAGGATTAAGAG
ATTAATTTCCCTAAAAATGATAAAACAAGCGTTTTGAAAGCGCTTGTTTTTTTGGTTTGCAGTCAGAGTAGAATAGAAG
TATCAAAAAAAGCACCGACTCGGTGCCACTTTTTCAAGTTGATAACGGACTAGCCTTATTTTAACTTGCTATGCTGTTT
TGAATGGTTCCAACAAGATTATTTTATAACTTTTATAACAAATAATCAAGGAGAAATTCAAAGAAATTTATCAGCCATA
AAACAATACTTAATACTATAGAATGATAACAAAATAAACTACTTTTTAAAAGAATTTTGTGTTATAATCTATTTATTAT
TAAGTATTGGGTAATATTTTTTGAAGAGATATTTTGAAAAAGAAAAATTAAAGCATATTAAACTAATTTCGGAGGTCAT
TAAAACTATTATTGAAATCATCAAACTCATTATGGATTTAATTTAAACTTTTTATTTTAGGAGGCAAAA**ATG**GATAAGA
AATACTCAATAGGCTTA**gct**ATCGGCACAAATAGCGTCGGATGGGCGGTGATCACTGATGAATATAAGGTTCCGTCTAA
AAAGTTCAAGGTTCTGGGAAATACAGACCGCCACAGTATCAAAAAAAATCTTATAGGGGCTCTTTTATTTGACAGTGGA
GAGACAGCGGAAGCGACTCGTCTCAAACGGACAGCTCGTAGAAGGTATACACGTCGGAAGAATCGTATTTGTTATCTAC
AGGAGATTTTTTCAAATGAGATGGCGAAAGTAGATGATAGTTTCTTTCATCGACTTGAAGAGTCTTTTTTGGTGGAAGA
AGACAAGAAGCATGAACGTCATCCTATTTTTGGAAATATAGTAGATGAAGTTGCTTATCATGAGAAATATCCAACTATC
TATCATCTGCGAAAAAAATTGGTAGATTCTACTGATAAAGCGGATTTGCGCTTAATCTATTTGGCCTTAGCGCATATGA
TTAAGTTTCGTGGTCATTTTTTGATTGAGGGAGATTTAAATCCTGATAATAGTGATGTGGACAAACTATTTATCCAGTT
GGTACAAACCTACAATCAATTATTTGAAGAAAACCCTATTAACGCAAGTGGAGTAGATGCTAAAGCGATTCTTTCTGCA
CGATTGAGTAAATCAAGACGATTAGAAAATCTCATTGCTCAGCTCCCCGGTGAGAAGAAAAATGGCTTATTTGGGAATC
TCATTGCTTTGTCATTGGGTTTGACCCCTAATTTTAAATCAAATTTTGATTTGGCAGAAGATGCTAAATTACAGCTTTC
AAAAGATACTTACGATGATGATTTAGATAATTTATTGGCGCAAATTGGAGATCAATATGCTGATTTGTTTTTGGCAGCT
AAGAATTTATCAGATGCTATTTTACTTTCAGATATCCTAAGAGTAAATACTGAAATAACTAAGGCTCCCCTATCAGCTT
CAATGATTAAACGCTACGATGAACATCATCAAGACTTGACTCTTTTAAAAGCTTTAGTTCGACAACAACTTCCAGAAAA
GTATAAAGAAATCTTTTTTGATCAATCAAAAAACGGATATGCAGGTTATATTGATGGGGGAGCTAGCCAAGAAGAATTT
TATAAATTTATCAAACCAATTTTAGAAAAAATGGATGGTACTGAGGAATTATTGGTGAAACTAAATCGTGAAGATTTGC
TGCGCAAGCAACGGACCTTTGACAACGGCTCTATTCCCCATCAAATTCACTTGGGTGAGCTGCATGCTATTTTGAGAAG
ACAAGAAGACTTTTATCCATTTTTAAAAGACAATCGTGAGAAGATTGAAAAAATCTTGACTTTTCGAATTCCTTATTAT
GTTGGTCCATTGGCGCGTGGCAATAGTCGTTTTGCATGGATGACTCGGAAGTCTGAAGAAACAATTACCCCATGGAATT
TTGAAGAAGTTGTCGATAAAGGTGCTTCAGCTCAATCATTTATTGAACGCATGACAAACTTTGATAAAAATCTTCCAAA
TGAAAAAGTACTACCAAAACATAGTTTGCTTTATGAGTATTTTACGGTTTATAACGAATTGACAAAGGTCAAATATGTT
ACTGAAGGAATGCGAAAACCAGCATTTCTTTCAGGTGAACAGAAGAAAGCCATTGTTGATTTACTCTTCAAAACAAATC
GAAAAGTAACCGTTAAGCAATTAAAAGAAGATTATTTCAAAAAAATAGAATGTTTTGATAGTGTTGAAATTTCAGGAGT
TGAAGATAGATTTAATGCTTCATTAGGTACCTACCATGATTTGCTAAAAATTATTAAAGATAAAGATTTTTTGGATAAT
GAAGAAAATGAAGATATCTTAGAGGATATTGTTTTAACATTGACCTTATTTGAAGATAGGGAGATGATTGAGGAAAGAC
TTAAAACATATGCTCACCTCTTTGATGATAAGGTGATGAAACAGCTTAAACGTCGCCGTTATACTGGTTGGGGACGTTT
GTCTCGAAAATTGATTAATGGTATTAGGGATAAGCAATCTGGCAAAACAATATTAGATTTTTTGAAATCAGATGGTTTT
GCCAATCGCAATTTTATGCAGCTGATCCATGATGATAGTTTGACATTTAAAGAAGACATTCAAAAAGCACAAGTGTCTG
GACAAGGCGATAGTTTACATGAACATATTGCAAATTTAGCTGGTAGCCCTGCTATTAAAAAAGGTATTTTACAGACTGT
AAAAGTTGTTGATGAATTGGTCAAAGTAATGGGGCGGCATAAGCCAGAAAATATCGTTATTGAAATGGCACGTGAAAAT
CAGACAACTCAAAAGGGCCAGAAAAATTCGCGAGAGCGTATGAAACGAATCGAAGAAGGTATCAAAGAATTAGGAAGTC
AGATTCTTAAAGAGCATCCTGTTGAAAATACTCAATTGCAAAATGAAAAGCTCTATCTCTATTATCTCCAAAATGGAAG
AGACATGTATGTGGACCAAGAATTAGATATTAATCGTTTAAGTGATTATGATGTCGAT**gcg**ATTGTTCCACAAAGTTTC
CTTAAAGACGATTCAATAGACAATAAGGTCTTAACGCGTTCTGATAAAAATCGTGGTAAATCGGATAACGTTCCAAGTG
AAGAAGTAGTCAAAAAGATGAAAAACTATTGGAGACAACTTCTAAACGCCAAGTTAATCACTCAACGTAAGTTTGATAA
TTTAACGAAAGCTGAACGTGGAGGTTTGAGTGAACTTGATAAAGCTGGTTTTATCAAACGCCAATTGGTTGAAACTCGC
CAAATCACTAAGCATGTGGCACAAATTTTGGATAGTCGCATGAATACTAAATACGATGAAAATGATAAACTTATTCGAG
AGGTTAAAGTGATTACCTTAAAATCTAAATTAGTTTCTGACTTCCGAAAAGATTTCCAATTCTATAAAGTACGTGAGAT
TAACAATTACCATCATGCCCATGATGCGTATCTAAATGCCGTCGTTGGAACTGCTTTGATTAAGAAATATCCAAAACTT
GAATCGGAGTTTGTCTATGGTGATTATAAAGTTTATGATGTTCGTAAAATGATTGCTAAGTCTGAGCAAGAAATAGGCA
AAGCAACCGCAAAATATTTCTTTTACTCTAATATCATGAACTTCTTCAAAACAGAAATTACACTTGCAAATGGAGAGAT
TCGCAAACGCCCTCTAATCGAAACTAATGGGGAAACTGGAGAAATTGTCTGGGATAAAGGGCGAGATTTTGCCACAGTG
CGCAAAGTATTGTCCATGCCCCAAGTCAATATTGTCAAGAAAACAGAAGTACAGACAGGCGGATTCTCCAAGGAGTCAA
TTTTACCAAAAAGAAATTCGGACAAGCTTATTGCTCGTAAAAAAGACTGGGATCCAAAAAAATATGGTGGTTTTGATAG
TCCAACGGTAGCTTATTCAGTCCTAGTGGTTGCTAAGGTGGAAAAAGGGAAATCGAAGAAGTTAAAATCCGTTAAAGAG
TTACTAGGGATCACAATTATGGAAAGAAGTTCCTTTGAAAAAAATCCGATTGACTTTTTAGAAGCTAAAGGATATAAGG
AAGTTAAAAAAGACTTAATCATTAAACTACCTAAATATAGTCTTTTTGAGTTAGAAAACGGTCGTAAACGGATGCTGGC
TAGTGCCGGAGAATTACAAAAAGGAAATGAGCTGGCTCTGCCAAGCAAATATGTGAATTTTTTATATTTAGCTAGTCAT

TATGAAAAGTTGAAGGGTAGTCCAGAAGATAACGAACAAAAACAATTGTTTGTGGAGCAGCATAAGCATTATTTAGATG
AGATTATTGAGCAAATCAGTGAATTTTCTAAGCGTGTTATTTTAGCAGATGCCAATTTAGATAAAGTTCTTAGTGCATA
TAACAAACATAGAGACAAACCAATACGTGAACAAGCAGAAAATATTATTCATTTATTTACGTTGACGAATCTTGGAGCT
CCCGCTGCTTTTAAATATTTTGATACAACAATTGATCGTAAACGATATACGTCTACAAAAGAAGTTTTAGATGCCACTC
TTATCCATCAATCCATCACTGGTCTTTATGAAACACGCATTGATTTGAGTCAGCTAGGAGGTGAC**TAA**CTCGAGTAAGG
ATCTCCAGGCATCAAATAAAACGAAAGGCTCAGTCGAAAGACTGGGCCTTTCGTTTTATCTGTTGTTTGTCGGTGAACG
CTCTCTACTAGAGTCACACTGGCTCACCTTCGGGTGGGCCTTTCTGCGTTTATA

>dxCas9(3.7)

**ATG**GACAAGAAGTACTCCATTGGGCTC**gct**ATCGGCACAAACAGCGTCGGCTGGGCCGTCATTACGGACGAGTACAAGG
TGCCGAGCAAAAAATTCAAAGTTCTGGGCAATACCGATCGCCACAGCATAAAGAAGAACCTCATTGGCGCCCTCCTGTT
CGACTCCGGGGAGACGGCCGAAGCCACGCGGCTCAAAAGAACAGCACGGCGCAGATATACCCGCAGAAAGAATCGGATC
TGCTACCTGCAGGAGATCTTTAGTAATGAGATGGCTAAGGTGGATGACTCTTTCTTCCATAGGCTGGAGGAGTCCTTTT
TGGTGGAGGAGGATAAAAAGCACGAGCGCCACCCAATCTTTGGCAATATCGTGGACGAGGTGGCGTACCATGAAAAGTA
CCCAACCATATATCATCTGAGGAAGAAGCTTGTAGACAGTACTGATAAGGCTGACTTGCGGTTGATCTATCTCGCGCTG
GCGCATATGATCAAATTTCGGGGACACTTCCTCATCGAGGGGGACCTGAACCCAGACAACAGCGATGTCGACAAACTCT
TTATCCAACTGGTTCAGACTTACAATCAGCTTTTCGAAGAGAACCCGATCAACGCATCCGGAGTTGACGCCAAAGCAAT
CCTGAGCGCTAGGCTGTCCAAATCCCGGCGGCTCGAAAACCTCATCGCACAGCTCCCTGGGGAGAAGAAGAACGGCCTG
TTTGGTAATCTTATCGCCCTGTCCCTCGGGCTGACCCCCAACTTTAAATCTAACTTCGACCTGGCCGAAGAT**acc**AAGC
TTCAACTGAGCAAAGACACCTACGATGATGATCTCGACAATCTGCTGGCCCAGATCGGCGACCAGTACGCAGACCTTTT
TTTGGCGGCAAAGAACCTGTCAGACGCCATTCTGCTGAGTGATATTCTGCGAGTGAACACGGAGATCACCAAAGCTCCG
CTGAGCGCTAGTATGATCAAG**ctc**TATGATGAGCACCACCAAGACTTGACTTTGCTGAAGGCCCTTGTCAGACAGCAAC
TGCCTGAGAAGTACAAGGAAATTTTCTTCGATCAGTCTAAAAATGGCTACGCCGGATACATTGACGGCGGAGCAAGCCA
GGAGGAATTTTACAAATTTATTAAGCCCATCTTGGAAAAAATGGACGGCACCGAGGAGCTGCTGGTAAAGCTTAACAGA
GAAGATCTGTTGCGCAAACAGCGCACTTTCGACAATGGA**atc**ATCCCCCACCAGATTCACCTGGGCGAACTGCACGCTA
TCCTCAGGCGGCAAGAGGATTTCTACCCCTTTTTGAAAGATAACAGGGAAAAGATTGAGAAAATCCTCACATTTCGGAT
ACCCTACTATGTAGGCCCCCTCGCCCGGGGAAATTCCAGATTCGCGTGGATGACTCGCAAATCAGAAGAGACCATCACT
CCCTGGAACTTCGAG**aaa**GTCGTGGATAAGGGGGCCTCTGCCCAGTCCTTCATCGAAAGGATGACTAACTTTGATAAAA
ATCTGCCTAACGAAAAGGTGCTTCCTAAACACTCTCTGCTGTACGAGTACTTCACAGTTTATAACGAGCTCACCAAGGT
CAAATACGTCACAGAAGGGATGAGAAAGCCAGCATTCCTGTCTGGA**gat**CAGAAGAAAGCTATTGTGGACCTCCTCTTC
AAGACGAACCGGAAAGTTACCGTGAAACAGCTCAAAGAAGACTATTTCAAAAAGATTGAATGTTTCGACTCTGTTGAAA
TCAGCGGAGTGGAGGATCGCTTCAACGCATCCCTGGGAACGTATCACGATCTCCTGAAAATCATTAAAGACAAGGACTT
CCTGGACAATGAGGAGAACGAGGACATTCTTGAGGACATTGTCCTCACCCTTACGTTGTTTGAAGATAGGGAGATGATT
GAAGAACGCTTGAAAACTTACGCTCATCTCTTCGACGACAAAGTCATGAAGCAGCTCAAGAGGCGCCGATATACAGGAT
GGGGGCGGCTGTCAAGAAAACTGATCAATGGGATCCGAGACAAGCAGAGTGGAAAGACAATCCTGGATTTTCTTAAGTC
CGATGGATTTGCCAACCGGAACTTC**att**CAGTTGATCCATGATGACTCTCTCACCTTTAAGGAGGACATCCAGAAAGCA
CAAGTTTCTGGCCAGGGGGACAGTCTTCACGAGCACATCGCTAATCTTGCAGGTAGCCCAGCTATCAAAAAGGGAATAC
TGCAGACCGTTAAGGTCGTGGATGAACTCGTCAAAGTAATGGGAAGGCATAAGCCCGAGAATATCGTTATCGAGATGGC
CCGAGAGAACCAAACCACCCAGAAGGGACAGAAGAACAGTAGGGAAAGGATGAAGAGGATTGAAGAGGGTATAAAAGAA
CTGGGGTCCCAAATCCTTAAGGAACACCCAGTTGAAAACACCCAGCTTCAGAATGAGAAGCTCTACCTGTACTACCTGC
AGAACGGCAGGGACATGTACGTGGATCAGGAACTGGACATCAATCGGCTCTCCGACTACGACGTGGAC**gct**ATCGTGCC
CCAGTCTTTTCTCAAAGATGATTCTATTGATAATAAAGTGTTGACAAGATCCGATAAAAACAGAGGGAAGAGTGATAAC
GTCCCCTCAGAAGAAGTTGTCAAGAAAATGAAAAATTATTGGCGGCAGCTGCTGAACGCCAAACTGATCACACAACGGA
AGTTCGATAATCTGACTAAGGCTGAACGAGGTGGCCTGTCTGAGTTGGATAAAGCCGGTTTCATCAAAAGGCAGCTTGT
TGAGACACGCCAGATCACCAAGCACGTGGCCCAAATTCTCGATTCACGCATGAACACCAAGTACGATGAAAATGACAAA
CTGATTCGAGAGGTGAAAGTTATTACTCTGAAGTCTAAGCTGGTCTCAGATTTCAGAAAGGACTTTCAGTTTTATAAGG
TGAGAGAGATCAACAATTACCACCATGCGCATGATGCCTACCTGAATGCAGTGGTAGGCACTGCACTTATCAAAAAATA
TCCCAAGCTTGAATCTGAATTTGTTTACGGAGACTATAAAGTGTACGATGTTAGGAAAATGATCGCAAAGTCTGAGCAG
GAAATAGGCAAGGCCACCGCTAAGTACTTCTTTTACAGCAATATTATGAATTTTTTCAAGACCGAGATTACACTGGCCA
ATGGAGAGATTCGGAAGCGACCACTTATCGAAACAAACGGAGAAACAGGAGAAATCGTGTGGGACAAGGGTAGGGATTT
CGCGACAGTCCGGAAGGTCCTGTCCATGCCGCAGGTGAACATCGTTAAAAAGACCGAAGTACAGACCGGAGGTTTCTCC
AAGGAAAGTATCCTCCCGAAAAGGAACAGCGACAAGCTGATCGCACGCAAAAAAGATTGGGACCCCAAGAAATACGGCG
GATTCGATTCTCCTACAGTCGCTTACAGTGTACTGGTTGTGGCCAAAGTGGAGAAAGGGAAGTCTAAAAAACTCAAAAG
CGTCAAGGAACTGCTGGGCATCACAATCATGGAGCGATCAAGCTTCGAAAAAAACCCCATCGACTTTCTCGAGGCGAAA
GGATATAAAGAGGTCAAAAAAGACCTCATCATTAAGCTTCCCAAGTACTCTCTCTTTGAGCTTGAAAACGGCCGGAAAC
GAATGCTCGCTAGTGCGGGC**gtg**CTGCAGAAAGGTAACGAGCTGGCACTGCCCTCTAAATACGTTAATTTCTTGTATCT

GGCCAGCCACTATGAAAAGCTCAAAGGGTCTCCCGAAGATAATGAGCAGAAGCAGCTGTTCGTGGAACAACACAAACAC
TACCTTGATGAGATCATCGAGCAAATAAGCGAATTCTCCAAAAGAGTGATCCTCGCCGACGCTAACCTCGATAAGGTGC
TTTCTGCTTACAATAAGCACAGGGATAAGCCCATCAGGGAGCAGGCAGAAAACATTATCCACTTGTTTACTCTGACCAA
CTTGGGCGCGCCTGCAGCCTTCAAGTACTTCGACACTACCATAGACAGAAAGCGGTACACCTCTACAAAGGAGGTCCTG
GACGCCACACTGATTCATCAGTCAATTACGGGGCTCTATGAAACAAGAATCGACCTCTCTCAGCTCGGTGGAGAC**TAA**

>dCas9-NG

**ATG**GATAAGAAATACTCAATAGGCTTA**gct**ATCGGCACAAATAGCGTCGGATGGGCGGTGATCACTGATGAATATAAGG
TTCCGTCTAAAAAGTTCAAGGTTCTGGGAAATACAGACCGCCACAGTATCAAAAAAAATCTTATAGGGGCTCTTTTATT
TGACAGTGGAGAGACAGCGGAAGCGACTCGTCTCAAACGGACAGCTCGTAGAAGGTATACACGTCGGAAGAATCGTATT
TGTTATCTACAGGAGATTTTTTCAAATGAGATGGCGAAAGTAGATGATAGTTTCTTTCATCGACTTGAAGAGTCTTTTT
TGGTGGAAGAAGACAAGAAGCATGAACGTCATCCTATTTTTGGAAATATAGTAGATGAAGTTGCTTATCATGAGAAATA
TCCAACTATCTATCATCTGCGAAAAAAATTGGTAGATTCTACTGATAAAGCGGATTTGCGCTTAATCTATTTGGCCTTA
GCGCATATGATTAAGTTTCGTGGTCATTTTTTGATTGAGGGAGATTTAAATCCTGATAATAGTGATGTGGACAAACTAT
TTATCCAGTTGGTACAAACCTACAATCAATTATTTGAAGAAAACCCTATTAACGCAAGTGGAGTAGATGCTAAAGCGAT
TCTTTCTGCACGATTGAGTAAATCAAGACGATTAGAAAATCTCATTGCTCAGCTCCCCGGTGAGAAGAAAAATGGCTTA
TTTGGGAATCTCATTGCTTTGTCATTGGGTTTGACCCCTAATTTTAAATCAAATTTTGATTTGGCAGAAGATGCTAAAT
TACAGCTTTCAAAAGATACTTACGATGATGATTTAGATAATTTATTGGCGCAAATTGGAGATCAATATGCTGATTTGTT
TTTGGCAGCTAAGAATTTATCAGATGCTATTTTACTTTCAGATATCCTAAGAGTAAATACTGAAATAACTAAGGCTCCC
CTATCAGCTTCAATGATTAAACGCTACGATGAACATCATCAAGACTTGACTCTTTTAAAAGCTTTAGTTCGACAACAAC
TTCCAGAAAAGTATAAAGAAATCTTTTTTGATCAATCAAAAAACGGATATGCAGGTTATATTGATGGGGGAGCTAGCCA
AGAAGAATTTTATAAATTTATCAAACCAATTTTAGAAAAAATGGATGGTACTGAGGAATTATTGGTGAAACTAAATCGT
GAAGATTTGCTGCGCAAGCAACGGACCTTTGACAACGGCTCTATTCCCCATCAAATTCACTTGGGTGAGCTGCATGCTA
TTTTGAGAAGACAAGAAGACTTTTATCCATTTTTAAAAGACAATCGTGAGAAGATTGAAAAAATCTTGACTTTTCGAAT
TCCTTATTATGTTGGTCCATTGGCGCGTGGCAATAGTCGTTTTGCATGGATGACTCGGAAGTCTGAAGAAACAATTACC
CCATGGAATTTTGAAGAAGTTGTCGATAAAGGTGCTTCAGCTCAATCATTTATTGAACGCATGACAAACTTTGATAAAA
ATCTTCCAAATGAAAAAGTACTACCAAAACATAGTTTGCTTTATGAGTATTTTACGGTTTATAACGAATTGACAAAGGT
CAAATATGTTACTGAAGGAATGCGAAAACCAGCATTTCTTTCAGGTGAACAGAAGAAAGCCATTGTTGATTTACTCTTC
AAAACAAATCGAAAAGTAACCGTTAAGCAATTAAAAGAAGATTATTTCAAAAAAATAGAATGTTTTGATAGTGTTGAAA
TTTCAGGAGTTGAAGATAGATTTAATGCTTCATTAGGTACCTACCATGATTTGCTAAAAATTATTAAAGATAAAGATTT
TTTGGATAATGAAGAAAATGAAGATATCTTAGAGGATATTGTTTTAACATTGACCTTATTTGAAGATAGGGAGATGATT
GAGGAAAGACTTAAAACATATGCTCACCTCTTTGATGATAAGGTGATGAAACAGCTTAAACGTCGCCGTTATACTGGTT
GGGGACGTTTGTCTCGAAAATTGATTAATGGTATTAGGGATAAGCAATCTGGCAAAACAATATTAGATTTTTTGAAATC
AGATGGTTTTGCCAATCGCAATTTTATGCAGCTGATCCATGATGATAGTTTGACATTTAAAGAAGACATTCAAAAAGCA
CAAGTGTCTGGACAAGGCGATAGTTTACATGAACATATTGCAAATTTAGCTGGTAGCCCTGCTATTAAAAAAGGTATTT
TACAGACTGTAAAAGTTGTTGATGAATTGGTCAAAGTAATGGGGCGGCATAAGCCAGAAAATATCGTTATTGAAATGGC
ACGTGAGAACCAAACCACCCAGAAGGGACAGAAGAACAGTAGGGAAAGGATGAAGAGGATTGAAGAGGGTATAAAAGAA
CTGGGGTCCCAAATCCTTAAGGAACACCCAGTTGAAAACACCCAGCTTCAGAATGAGAAGCTCTACCTGTACTACCTGC
AGAACGGCAGGGACATGTACGTGGATCAGGAACTGGACATCAATCGGCTCTCCGACTACGACGTGGAC**gct**ATCGTGCC
CCAGTCTTTTCTCAAAGATGATTCTATTGATAATAAAGTGTTGACAAGATCCGATAAAAACAGAGGGAAGAGTGATAAC
GTCCCCTCAGAAGAAGTTGTCAAGAAAATGAAAAATTATTGGCGGCAGCTGCTGAACGCCAAACTGATCACACAACGGA
AGTTCGATAATCTGACTAAGGCTGAACGAGGTGGCCTGTCTGAGTTGGATAAAGCCGGTTTCATCAAAAGGCAGCTTGT
TGAGACACGCCAGATCACCAAGCACGTGGCCCAAATTCTCGATTCACGCATGAACACCAAGTACGATGAAAATGACAAA
CTGATTCGAGAGGTGAAAGTTATTACTCTGAAGTCTAAGCTGGTCTCAGATTTCAGAAAGGACTTTCAGTTTTATAAGG
TGAGAGAGATCAACAATTACCACCATGCGCATGATGCCTACCTGAATGCAGTGGTAGGCACTGCACTTATCAAAAAATA
TCCCAAGCTTGAATCTGAATTTGTTTACGGAGACTATAAAGTGTACGATGTTAGGAAAATGATCGCAAAGTCTGAGCAG
GAAATAGGCAAGGCCACCGCTAAGTACTTCTTTTACAGCAATATTATGAATTTTTTCAAGACCGAGATTACACTGGCCA
ATGGAGAGATTCGGAAGCGACCACTTATCGAAACAAACGGAGAAACAGGAGAAATCGTGTGGGACAAGGGTAGGGATTT
CGCGACAGTCCGGAAGGTCCTGTCCATGCCGCAGGTGAACATCGTTAAAAAGACCGAAGTACAGACCGGAGGTTTCTCC
AAGGAAAGTATC**cgt**CCGAAAAGGAACAGCGACAAGCTGATCGCACGCAAAAAAGATTGGGACCCCAAGAAATACGGCG
GATTC**gtg**TCTCCTACAGTCGCTTACAGTGTACTGGTTGTGGCCAAAGTGGAGAAAGGGAAGTCTAAAAAACTCAAAAG
CGTCAAGGAACTGCTGGGCATCACAATCATGGAGCGATCAAGCTTCGAAAAAAACCCCATCGACTTTCTGGAGGCGAAA
GGATATAAAGAGGTCAAAAAAGACCTCATCATTAAGCTTCCCAAGTACTCTCTCTTTGAGCTTGAAAACGGCCGGAAAC
GAATGCTCGCTAGTGCG**cgtttt**CTGCAGAAAGGTAACGAGCTGGCACTGCCCTCTAAATACGTTAATTTCTTGTATCT
GGCCAGCCACTATGAAAAGCTCAAAGGGTCTCCCGAAGATAATGAGCAGAAGCAGCTGTTCGTGGAACAACACAAACAC
TACCTTGATGAGATCATCGAGCAAATAAGCGAATTCTCCAAAAGAGTGATCCTCGCCGACGCTAACCTCGATAAGGTGC

TTTCTGCTTACAATAAGCACAGGGATAAGCCCATCAGGGAGCAGGCAGAAAACATTATCCACTTGTTTACTCTGACCAA
CTTGGGCGCGCCT**cgt**GCCTTCAAGTACTTCGACACTACCATAGACAGAAAG**gtg**TAC**cgc**TCTACAAAGGAGGTCCTG
GACGCCACACTGATTCATCAGTCAATTACGGGGCTCTATGAAACAAGAATCGACCTCTCTCAGCTCGGTGGAGAC**TAA**

>dxCas9-NG

**ATG**GACAAGAAGTACTCCATTGGGCTC**gct**ATCGGCACAAACAGCGTCGGCTGGGCCGTCATTACGGACGAGTACAAGG
TGCCGAGCAAAAAATTCAAAGTTCTGGGCAATACCGATCGCCACAGCATAAAGAAGAACCTCATTGGCGCCCTCCTGTT
CGACTCCGGGGAGACGGCCGAAGCCACGCGGCTCAAAAGAACAGCACGGCGCAGATATACCCGCAGAAAGAATCGGATC
TGCTACCTGCAGGAGATCTTTAGTAATGAGATGGCTAAGGTGGATGACTCTTTCTTCCATAGGCTGGAGGAGTCCTTTT
TGGTGGAGGAGGATAAAAAGCACGAGCGCCACCCAATCTTTGGCAATATCGTGGACGAGGTGGCGTACCATGAAAAGTA
CCCAACCATATATCATCTGAGGAAGAAGCTTGTAGACAGTACTGATAAGGCTGACTTGCGGTTGATCTATCTCGCGCTG
GCGCATATGATCAAATTTCGGGGACACTTCCTCATCGAGGGGGACCTGAACCCAGACAACAGCGATGTCGACAAACTCT
TTATCCAACTGGTTCAGACTTACAATCAGCTTTTCGAAGAGAACCCGATCAACGCATCCGGAGTTGACGCCAAAGCAAT
CCTGAGCGCTAGGCTGTCCAAATCCCGGCGGCTCGAAAACCTCATCGCACAGCTCCCTGGGGAGAAGAAGAACGGCCTG
TTTGGTAATCTTATCGCCCTGTCCCTCGGGCTGACCCCCAACTTTAAATCTAACTTCGACCTGGCCGAAGAT**acc**AAGC
TTCAACTGAGCAAAGACACCTACGATGATGATCTCGACAATCTGCTGGCCCAGATCGGCGACCAGTACGCAGACCTTTT
TTTGGCGGCAAAGAACCTGTCAGACGCCATTCTGCTGAGTGATATTCTGCGAGTGAACACGGAGATCACCAAAGCTCCG
CTGAGCGCTAGTATGATCAAG**ctc**TATGATGAGCACCACCAAGACTTGACTTTGCTGAAGGCCCTTGTCAGACAGCAAC
TGCCTGAGAAGTACAAGGAAATTTTCTTCGATCAGTCTAAAAATGGCTACGCCGGATACATTGACGGCGGAGCAAGCCA
GGAGGAATTTTACAAATTTATTAAGCCCATCTTGGAAAAAATGGACGGCACCGAGGAGCTGCTGGTAAAGCTTAACAGA
GAAGATCTGTTGCGCAAACAGCGCACTTTCGACAATGGA**atc**ATCCCCCACCAGATTCACCTGGGCGAACTGCACGCTA
TCCTCAGGCGGCAAGAGGATTTCTACCCCTTTTTGAAAGATAACAGGGAAAAGATTGAGAAAATCCTCACATTTCGGAT
ACCCTACTATGTAGGCCCCCTCGCCCGGGGAAATTCCAGATTCGCGTGGATGACTCGCAAATCAGAAGAGACCATCACT
CCCTGGAACTTCGAG**aaa**GTCGTGGATAAGGGGGCCTCTGCCCAGTCCTTCATCGAAAGGATGACTAACTTTGATAAAA
ATCTGCCTAACGAAAAGGTGCTTCCTAAACACTCTCTGCTGTACGAGTACTTCACAGTTTATAACGAGCTCACCAAGGT
CAAATACGTCACAGAAGGGATGAGAAAGCCAGCATTCCTGTCTGGA**gat**CAGAAGAAAGCTATTGTGGACCTCCTCTTC
AAGACGAACCGGAAAGTTACCGTGAAACAGCTCAAAGAAGACTATTTCAAAAAGATTGAATGTTTCGACTCTGTTGAAA
TCAGCGGAGTGGAGGATCGCTTCAACGCATCCCTGGGAACGTATCACGATCTCCTGAAAATCATTAAAGACAAGGACTT
CCTGGACAATGAGGAGAACGAGGACATTCTTGAGGACATTGTCCTCACCCTTACGTTGTTTGAAGATAGGGAGATGATT
GAAGAACGCTTGAAAACTTACGCTCATCTCTTCGACGACAAAGTCATGAAGCAGCTCAAGAGGCGCCGATATACAGGAT
GGGGGCGGCTGTCAAGAAAACTGATCAATGGGATCCGAGACAAGCAGAGTGGAAAGACAATCCTGGATTTTCTTAAGTC
CGATGGATTTGCCAACCGGAACTTC**att**CAGTTGATCCATGATGACTCTCTCACCTTTAAGGAGGACATCCAGAAAGCA
CAAGTTTCTGGCCAGGGGGACAGTCTTCACGAGCACATCGCTAATCTTGCAGGTAGCCCAGCTATCAAAAAGGGAATAC
TGCAGACCGTTAAGGTCGTGGATGAACTCGTCAAAGTAATGGGAAGGCATAAGCCCGAGAATATCGTTATCGAGATGGC
CCGAGAGAACCAAACCACCCAGAAGGGACAGAAGAACAGTAGGGAAAGGATGAAGAGGATTGAAGAGGGTATAAAAGAA
CTGGGGTCCCAAATCCTTAAGGAACACCCAGTTGAAAACACCCAGCTTCAGAATGAGAAGCTCTACCTGTACTACCTGC
AGAACGGCAGGGACATGTACGTGGATCAGGAACTGGACATCAATCGGCTCTCCGACTACGACGTGGAC**gct**ATCGTGCC
CCAGTCTTTTCTCAAAGATGATTCTATTGATAATAAAGTGTTGACAAGATCCGATAAAAACAGAGGGAAGAGTGATAAC
GTCCCCTCAGAAGAAGTTGTCAAGAAAATGAAAAATTATTGGCGGCAGCTGCTGAACGCCAAACTGATCACACAACGGA
AGTTCGATAATCTGACTAAGGCTGAACGAGGTGGCCTGTCTGAGTTGGATAAAGCCGGTTTCATCAAAAGGCAGCTTGT
TGAGACACGCCAGATCACCAAGCACGTGGCCCAAATTCTCGATTCACGCATGAACACCAAGTACGATGAAAATGACAAA
CTGATTCGAGAGGTGAAAGTTATTACTCTGAAGTCTAAGCTGGTCTCAGATTTCAGAAAGGACTTTCAGTTTTATAAGG
TGAGAGAGATCAACAATTACCACCATGCGCATGATGCCTACCTGAATGCAGTGGTAGGCACTGCACTTATCAAAAAATA
TCCCAAGCTTGAATCTGAATTTGTTTACGGAGACTATAAAGTGTACGATGTTAGGAAAATGATCGCAAAGTCTGAGCAG
GAAATAGGCAAGGCCACCGCTAAGTACTTCTTTTACAGCAATATTATGAATTTTTTCAAGACCGAGATTACACTGGCCA
ATGGAGAGATTCGGAAGCGACCACTTATCGAAACAAACGGAGAAACAGGAGAAATCGTGTGGGACAAGGGTAGGGATTT
CGCGACAGTCCGGAAGGTCCTGTCCATGCCGCAGGTGAACATCGTTAAAAAGACCGAAGTACAGACCGGAGGTTTCTCC
AAGGAAAGTATC**cgt**CCGAAAAGGAACAGCGACAAGCTGATCGCACGCAAAAAAGATTGGGACCCCAAGAAATACGGCG
GATTC**gtg**TCTCCTACAGTCGCTTACAGTGTACTGGTTGTGGCCAAAGTGGAGAAAGGGAAGTCTAAAAAACTCAAAAG
CGTCAAGGAACTGCTGGGCATCACAATCATGGAGCGATCAAGCTTCGAAAAAAACCCCATCGACTTTCTGGAGGCGAAA
GGATATAAAGAGGTCAAAAAAGACCTCATCATTAAGCTTCCCAAGTACTCTCTCTTTGAGCTTGAAAACGGCCGGAAAC
GAATGCTCGCTAGTGCG**cgtttt**CTGCAGAAAGGTAACGAGCTGGCACTGCCCTCTAAATACGTTAATTTCTTGTATCT
GGCCAGCCACTATGAAAAGCTCAAAGGGTCTCCCGAAGATAATGAGCAGAAGCAGCTGTTCGTGGAACAACACAAACAC
TACCTTGATGAGATCATCGAGCAAATAAGCGAATTCTCCAAAAGAGTGATCCTCGCCGACGCTAACCTCGATAAGGTGC
TTTCTGCTTACAATAAGCACAGGGATAAGCCCATCAGGGAGCAGGCAGAAAACATTATCCACTTGTTTACTCTGACCAA

CTTGGGCGCGCCT**cgt**GCCTTCAAGTACTTCGACACTACCATAGACAGAAAG**gtg**TAC**cgc**TCTACAAAGGAGGTCCTG
GACGCCACACTGATTCATCAGTCAATTACGGGGCTCTATGAAACAAGAATCGACCTCTCTCAGCTCGGTGGAGAC**TAA**

>dSpG

**ATG**GATAAGAAATACTCAATAGGCTTA**gct**ATCGGCACAAATAGCGTCGGATGGGCGGTGATCACTGATGAATATAAGG
TTCCGTCTAAAAAGTTCAAGGTTCTGGGAAATACAGACCGCCACAGTATCAAAAAAAATCTTATAGGGGCTCTTTTATT
TGACAGTGGAGAGACAGCGGAAGCGACTCGTCTCAAACGGACAGCTCGTAGAAGGTATACACGTCGGAAGAATCGTATT
TGTTATCTACAGGAGATTTTTTCAAATGAGATGGCGAAAGTAGATGATAGTTTCTTTCATCGACTTGAAGAGTCTTTTT
TGGTGGAAGAAGACAAGAAGCATGAACGTCATCCTATTTTTGGAAATATAGTAGATGAAGTTGCTTATCATGAGAAATA
TCCAACTATCTATCATCTGCGAAAAAAATTGGTAGATTCTACTGATAAAGCGGATTTGCGCTTAATCTATTTGGCCTTA
GCGCATATGATTAAGTTTCGTGGTCATTTTTTGATTGAGGGAGATTTAAATCCTGATAATAGTGATGTGGACAAACTAT
TTATCCAGTTGGTACAAACCTACAATCAATTATTTGAAGAAAACCCTATTAACGCAAGTGGAGTAGATGCTAAAGCGAT
TCTTTCTGCACGATTGAGTAAATCAAGACGATTAGAAAATCTCATTGCTCAGCTCCCCGGTGAGAAGAAAAATGGCTTA
TTTGGGAATCTCATTGCTTTGTCATTGGGTTTGACCCCTAATTTTAAATCAAATTTTGATTTGGCAGAAGATGCTAAAT
TACAGCTTTCAAAAGATACTTACGATGATGATTTAGATAATTTATTGGCGCAAATTGGAGATCAATATGCTGATTTGTT
TTTGGCAGCTAAGAATTTATCAGATGCTATTTTACTTTCAGATATCCTAAGAGTAAATACTGAAATAACTAAGGCTCCC
CTATCAGCTTCAATGATTAAACGCTACGATGAACATCATCAAGACTTGACTCTTTTAAAAGCTTTAGTTCGACAACAAC
TTCCAGAAAAGTATAAAGAAATCTTTTTTGATCAATCAAAAAACGGATATGCAGGTTATATTGATGGGGGAGCTAGCCA
AGAAGAATTTTATAAATTTATCAAACCAATTTTAGAAAAAATGGATGGTACTGAGGAATTATTGGTGAAACTAAATCGT
GAAGATTTGCTGCGCAAGCAACGGACCTTTGACAACGGCTCTATTCCCCATCAAATTCACTTGGGTGAGCTGCATGCTA
TTTTGAGAAGACAAGAAGACTTTTATCCATTTTTAAAAGACAATCGTGAGAAGATTGAAAAAATCTTGACTTTTCGAAT
TCCTTATTATGTTGGTCCATTGGCGCGTGGCAATAGTCGTTTTGCATGGATGACTCGGAAGTCTGAAGAAACAATTACC
CCATGGAATTTTGAAGAAGTTGTCGATAAAGGTGCTTCAGCTCAATCATTTATTGAACGCATGACAAACTTTGATAAAA
ATCTTCCAAATGAAAAAGTACTACCAAAACATAGTTTGCTTTATGAGTATTTTACGGTTTATAACGAATTGACAAAGGT
CAAATATGTTACTGAAGGAATGCGAAAACCAGCATTTCTTTCAGGTGAACAGAAGAAAGCCATTGTTGATTTACTCTTC
AAAACAAATCGAAAAGTAACCGTTAAGCAATTAAAAGAAGATTATTTCAAAAAAATAGAATGTTTTGATAGTGTTGAAA
TTTCAGGAGTTGAAGATAGATTTAATGCTTCATTAGGTACCTACCATGATTTGCTAAAAATTATTAAAGATAAAGATTT
TTTGGATAATGAAGAAAATGAAGATATCTTAGAGGATATTGTTTTAACATTGACCTTATTTGAAGATAGGGAGATGATT
GAGGAAAGACTTAAAACATATGCTCACCTCTTTGATGATAAGGTGATGAAACAGCTTAAACGTCGCCGTTATACTGGTT
GGGGACGTTTGTCTCGAAAATTGATTAATGGTATTAGGGATAAGCAATCTGGCAAAACAATATTAGATTTTTTGAAATC
AGATGGTTTTGCCAATCGCAATTTTATGCAGCTGATCCATGATGATAGTTTGACATTTAAAGAAGACATTCAAAAAGCA
CAAGTGTCTGGACAAGGCGATAGTTTACATGAACATATTGCAAATTTAGCTGGTAGCCCTGCTATTAAAAAAGGTATTT
TACAGACTGTAAAAGTTGTTGATGAATTGGTCAAAGTAATGGGGCGGCATAAGCCAGAAAATATCGTTATTGAAATGGC
ACGTGAAAATCAGACAACTCAAAAGGGCCAGAAAAATTCGCGAGAGCGTATGAAACGAATCGAAGAAGGTATCAAAGAA
TTAGGAAGTCAGATTCTTAAAGAGCATCCTGTTGAAAATACTCAATTGCAAAATGAAAAGCTCTATCTCTATTATCTCC
AAAATGGAAGAGACATGTATGTGGACCAAGAATTAGATATTAATCGTTTAAGTGATTATGATGTCGAT**gcg**ATTGTTCC
ACAAAGTTTCCTTAAAGACGATTCAATAGACAATAAGGTCTTAACGCGTTCTGATAAAAATCGTGGTAAATCGGATAAC
GTTCCAAGTGAAGAAGTAGTCAAAAAGATGAAAAACTATTGGAGACAACTTCTAAACGCCAAGTTAATCACTCAACGTA
AGTTTGATAATTTAACGAAAGCTGAACGTGGAGGTTTGAGTGAACTTGATAAAGCTGGTTTTATCAAACGCCAATTGGT
TGAAACTCGCCAAATCACTAAGCATGTGGCACAAATTTTGGATAGTCGCATGAATACTAAATACGATGAAAATGATAAA
CTTATTCGAGAGGTTAAAGTGATTACCTTAAAATCTAAATTAGTTTCTGACTTCCGAAAAGATTTCCAATTCTATAAAG
TACGTGAGATTAACAATTACCATCATGCCCATGATGCGTATCTAAATGCCGTCGTTGGAACTGCTTTGATTAAGAAATA
TCCAAAACTTGAATCGGAGTTTGTCTATGGTGATTATAAAGTTTATGATGTTCGTAAAATGATTGCTAAGTCTGAGCAA
GAAATAGGCAAAGCAACCGCAAAATATTTCTTTTACTCTAATATCATGAACTTCTTCAAAACAGAAATTACACTTGCAA
ATGGAGAGATTCGCAAACGCCCTCTAATCGAAACTAATGGGGAAACTGGAGAAATTGTCTGGGATAAAGGGCGAGATTT
TGCCACAGTGCGCAAAGTATTGTCCATGCCCCAAGTCAATATTGTCAAGAAAACAGAAGTACAGACAGGCGGATTCTCC
AAGGAGTCAATTTTACCAAAAAGAAATTCGGACAAGCTTATTGCTCGTAAAAAAGACTGGGATCCAAAAAAATATGGTG
GTTTT**ctgtgg**CCAACGGTAGCTTATTCAGTCCTAGTGGTTGCTAAGGTGGAAAAAGGGAAATCGAAGAAGTTAAAATC
CGTTAAAGAGTTACTAGGGATCACAATTATGGAAAGAAGTTCCTTTGAAAAAAATCCGATTGACTTTTTAGAAGCTAAA
GGATATAAGGAAGTTAAAAAAGACTTAATCATTAAACTACCTAAATATAGTCTTTTTGAGTTAGAAAACGGTCGTAAAC
GGATGCTGGCTAGTGCC**aaacag**TTACAAAAAGGAAATGAGCTGGCTCTGCCAAGCAAATATGTGAATTTTTTATATTT
AGCTAGTCATTATGAAAAGTTGAAGGGTAGTCCAGAAGATAACGAACAAAAACAATTGTTTGTGGAGCAGCATAAGCAT
TATTTAGATGAGATTATTGAGCAAATCAGTGAATTTTCTAAGCGTGTTATTTTAGCAGATGCCAATTTAGATAAAGTTC
TTAGTGCATATAACAAACATAGAGACAAACCAATACGTGAACAAGCAGAAAATATTATTCATTTATTTACGTTGACGAA
TCTTGGAGCTCCCGCTGCTTTTAAATATTTTGATACAACAATTGATCGTAAA**cag**TAT**cgt**TCTACAAAAGAAGTTTTA
GATGCCACTCTTATCCATCAATCCATCACTGGTCTTTATGAAACACGCATTGATTTGAGTCAGCTAGGAGGTGAC**TAA**

>dSpRY

**ATG**GATAAGAAATACTCAATAGGCTTA**gct**ATCGGCACAAATAGCGTCGGATGGGCGGTGATCACTGATGAATATAAGG
TTCCGTCTAAAAAGTTCAAGGTTCTGGGAAATACAGACCGCCACAGTATCAAAAAAAATCTTATAGGGGCTCTTTTATT
TGACAGTGGAGAGACAGCGGAA**cgt**ACTCGTCTCAAACGGACAGCTCGTAGAAGGTATACACGTCGGAAGAATCGTATT
TGTTATCTACAGGAGATTTTTTCAAATGAGATGGCGAAAGTAGATGATAGTTTCTTTCATCGACTTGAAGAGTCTTTTT
TGGTGGAAGAAGACAAGAAGCATGAACGTCATCCTATTTTTGGAAATATAGTAGATGAAGTTGCTTATCATGAGAAATA
TCCAACTATCTATCATCTGCGAAAAAAATTGGTAGATTCTACTGATAAAGCGGATTTGCGCTTAATCTATTTGGCCTTA
GCGCATATGATTAAGTTTCGTGGTCATTTTTTGATTGAGGGAGATTTAAATCCTGATAATAGTGATGTGGACAAACTAT
TTATCCAGTTGGTACAAACCTACAATCAATTATTTGAAGAAAACCCTATTAACGCAAGTGGAGTAGATGCTAAAGCGAT
TCTTTCTGCACGATTGAGTAAATCAAGACGATTAGAAAATCTCATTGCTCAGCTCCCCGGTGAGAAGAAAAATGGCTTA
TTTGGGAATCTCATTGCTTTGTCATTGGGTTTGACCCCTAATTTTAAATCAAATTTTGATTTGGCAGAAGATGCTAAAT
TACAGCTTTCAAAAGATACTTACGATGATGATTTAGATAATTTATTGGCGCAAATTGGAGATCAATATGCTGATTTGTT
TTTGGCAGCTAAGAATTTATCAGATGCTATTTTACTTTCAGATATCCTAAGAGTAAATACTGAAATAACTAAGGCTCCC
CTATCAGCTTCAATGATTAAACGCTACGATGAACATCATCAAGACTTGACTCTTTTAAAAGCTTTAGTTCGACAACAAC
TTCCAGAAAAGTATAAAGAAATCTTTTTTGATCAATCAAAAAACGGATATGCAGGTTATATTGATGGGGGAGCTAGCCA
AGAAGAATTTTATAAATTTATCAAACCAATTTTAGAAAAAATGGATGGTACTGAGGAATTATTGGTGAAACTAAATCGT
GAAGATTTGCTGCGCAAGCAACGGACCTTTGACAACGGCTCTATTCCCCATCAAATTCACTTGGGTGAGCTGCATGCTA
TTTTGAGAAGACAAGAAGACTTTTATCCATTTTTAAAAGACAATCGTGAGAAGATTGAAAAAATCTTGACTTTTCGAAT
TCCTTATTATGTTGGTCCATTGGCGCGTGGCAATAGTCGTTTTGCATGGATGACTCGGAAGTCTGAAGAAACAATTACC
CCATGGAATTTTGAAGAAGTTGTCGATAAAGGTGCTTCAGCTCAATCATTTATTGAACGCATGACAAACTTTGATAAAA
ATCTTCCAAATGAAAAAGTACTACCAAAACATAGTTTGCTTTATGAGTATTTTACGGTTTATAACGAATTGACAAAGGT
CAAATATGTTACTGAAGGAATGCGAAAACCAGCATTTCTTTCAGGTGAACAGAAGAAAGCCATTGTTGATTTACTCTTC
AAAACAAATCGAAAAGTAACCGTTAAGCAATTAAAAGAAGATTATTTCAAAAAAATAGAATGTTTTGATAGTGTTGAAA
TTTCAGGAGTTGAAGATAGATTTAATGCTTCATTAGGTACCTACCATGATTTGCTAAAAATTATTAAAGATAAAGATTT
TTTGGATAATGAAGAAAATGAAGATATCTTAGAGGATATTGTTTTAACATTGACCTTATTTGAAGATAGGGAGATGATT
GAGGAAAGACTTAAAACATATGCTCACCTCTTTGATGATAAGGTGATGAAACAGCTTAAACGTCGCCGTTATACTGGTT
GGGGACGTTTGTCTCGAAAATTGATTAATGGTATTAGGGATAAGCAATCTGGCAAAACAATATTAGATTTTTTGAAATC
AGATGGTTTTGCCAATCGCAATTTTATGCAGCTGATCCATGATGATAGTTTGACATTTAAAGAAGACATTCAAAAAGCA
CAAGTGTCTGGACAAGGCGATAGTTTACATGAACATATTGCAAATTTAGCTGGTAGCCCTGCTATTAAAAAAGGTATTT
TACAGACTGTAAAAGTTGTTGATGAATTGGTCAAAGTAATGGGGCGGCATAAGCCAGAAAATATCGTTATTGAAATGGC
ACGTGAAAATCAGACAACTCAAAAGGGCCAGAAAAATTCGCGAGAGCGTATGAAACGAATCGAAGAAGGTATCAAAGAA
TTAGGAAGTCAGATTCTTAAAGAGCATCCTGTTGAAAATACTCAATTGCAAAATGAAAAGCTCTATCTCTATTATCTCC
AAAATGGAAGAGACATGTATGTGGACCAAGAATTAGATATTAATCGTTTAAGTGATTATGATGTCGAT**gcg**ATTGTTCC
ACAAAGTTTCCTTAAAGACGATTCAATAGACAATAAGGTCTTAACGCGTTCTGATAAAAATCGTGGTAAATCGGATAAC
GTTCCAAGTGAAGAAGTAGTCAAAAAGATGAAAAACTATTGGAGACAACTTCTAAACGCCAAGTTAATCACTCAACGTA
AGTTTGATAATTTAACGAAAGCTGAACGTGGAGGTTTGAGTGAACTTGATAAAGCTGGTTTTATCAAACGCCAATTGGT
TGAAACTCGCCAAATCACTAAGCATGTGGCACAAATTTTGGATAGTCGCATGAATACTAAATACGATGAAAATGATAAA
CTTATTCGAGAGGTTAAAGTGATTACCTTAAAATCTAAATTAGTTTCTGACTTCCGAAAAGATTTCCAATTCTATAAAG
TACGTGAGATTAACAATTACCATCATGCCCATGATGCGTATCTAAATGCCGTCGTTGGAACTGCTTTGATTAAGAAATA
TCCAAAACTTGAATCGGAGTTTGTCTATGGTGATTATAAAGTTTATGATGTTCGTAAAATGATTGCTAAGTCTGAGCAA
GAAATAGGCAAAGCAACCGCAAAATATTTCTTTTACTCTAATATCATGAACTTCTTCAAAACAGAAATTACACTTGCAA
ATGGAGAGATTCGCAAACGCCCTCTAATCGAAACTAATGGGGAAACTGGAGAAATTGTCTGGGATAAAGGGCGAGATTT
TGCCACAGTGCGCAAAGTATTGTCCATGCCCCAAGTCAATATTGTCAAGAAAACAGAAGTACAGACAGGCGGATTCTCC
AAGGAGTCAATT**cgt**CCAAAAAGAAATTCGGACAAGCTTATTGCTCGTAAAAAAGACTGGGATCCAAAAAAATATGGTG
GTTTT**ctgtgg**CCAACGGTAGCTTATTCAGTCCTAGTGGTTGCTAAGGTGGAAAAAGGGAAATCGAAGAAGTTAAAATC
CGTTAAAGAGTTACTAGGGATCACAATTATGGAAAGAAGTTCCTTTGAAAAAAATCCGATTGACTTTTTAGAAGCTAAA
GGATATAAGGAAGTTAAAAAAGACTTAATCATTAAACTACCTAAATATAGTCTTTTTGAGTTAGAAAACGGTCGTAAAC
GGATGCTGGCTAGTGCC**aaacag**TTACAAAAAGGAAATGAGCTGGCTCTGCCAAGCAAATATGTGAATTTTTTATATTT
AGCTAGTCATTATGAAAAGTTGAAGGGTAGTCCAGAAGATAACGAACAAAAACAATTGTTTGTGGAGCAGCATAAGCAT
TATTTAGATGAGATTATTGAGCAAATCAGTGAATTTTCTAAGCGTGTTATTTTAGCAGATGCCAATTTAGATAAAGTTC
TTAGTGCATATAACAAACATAGAGACAAACCAATACGTGAACAAGCAGAAAATATTATTCATTTATTTACGTTGACG**cg**
**t**CTTGGAGCTCCC**cgt**GCTTTTAAATATTTTGATACAACAATTGATccgAAA**cag**TAT**cgt**TCTACAAAAGAAGTTTTA
GATGCCACTCTTATCCATCAATCCATCACTGGTCTTTATGAAACACGCATTGATTTGAGTCAGCTAGGAGGTGAC**TAA**

scRNA/sgRNA expression cassettes

>BBa_J23119(SpeI)-scRNA.1xMS2.b2-TrrnB

TTGACAGCTAGCTCAGTCCTAGGTATAATACTAGTNNNNNNNNNNNNNNNNNNNNGTTTTAGAGCTAGAAATAGCAAGT
TAAAATAAGGCTAGTCCGTTATCAACTTGAAAAAGTGGCACATGAGGATCACCCATGTGCTTTTTTTGAAGCTTGGGCC
CGAACAAAAACTCATCTCAGAAGAGGATCTGAATAGCGCCGTCGACCATCATCATCATCATCATTGAGTTTAAACGGTC
TCCAGCTTGGCTGTTTTGGCGGATGAGAGAAGATTTTCAGCCTGATACAGATTAAATCAGAACGCAGAAGCGGTCTGAT
AAAACAGAATTTGCCTGGCGGCAGTAGCGCGGTGGTCCCACCTGACCCCATGCCGAACTCAGAAGTGAAACGCCGTAGC
GCCGATGGTAGTGTGGGGTCTCCCCATGCGAGAGTAGGGAACTGCCAGGCATCAAATAAAACGAAAGGCTCAGTCGAAA
GACTGGGCCTTTCGTTTTATCTGTTGTTTGTCGGTGAACT

>BBa_J23119(SpeI)-sgRNA-BBa_K2680405

TTGACAGCTAGCTCAGTCCTAGGTATAATACTAGTNNNNNNNNNNNNNNNNNNNNGTTTTAGAGCTAGAAATAGCAAGT
TAAAATAAGGCTAGTCCGTTATCAACTTGAAAAAGTGGCACCGAGTCGGTGCTTTTTTTAACGCATGAGAAAGCCCCCG
GAAGATCACCTTCCGGGGGCTTTTTTATTGCGC

*E. coli* endogenous promoters (annotated by RegulonDB) as in pCK590 ( + strand) and pCK591 ( – strand). Underlined sequences are 35bp minimal promoters. **Bolded** nucleotides are **TSS** according to RegulonDB (cite Santos-Zavaleta et al., 2019) or **start codon** of the GFPmut2.

>yajGp
GGATCCGAACGCCGCCCTTAATTTTTCTTCTATCCGCTCACGTATCATCATGAAATATCTCCTCCGACAACGCTGAGAT
GTCATCCATCCCTTTAAATACTAGCCGCTTTTACCGTTTCCATTGCAGCTTAACAACAAATATTTAGGTTTCGTCCTGG
TTTTTCCGTCAAGCGTATGAAGTTTACTGATTTAAAGATTGAAACACAGGCCTCGCATCAGTGTTTCTTTTACCAGGGG
CGATTATTTCGTCAACATGATGAATTTACATGCCTTACCCACTTCCCCTCGCCGTTGGCAATGTTATGATGGCGG**G**AAT
TTTTACCACTCGAGAGATCCTCTAGATTTAAGAAGGAGATATACAT**ATG**

>uxuRp
CTCGAGACTTTGTTGGCGCAGTGACGGCGGCATATCAGCAGCTGTGCGAACGCGGTGCGCGCGAGTGTGTGGCTGCGCT
GTAACTAACTGATTACCCTACAGACTTACTGGTCAATCAAACTGATATTTGGTTGACCAGTTTTCGTTTTTTTGCCCAC
CTGTACGTGCCAACTTCCAGT**G**CTAATGGTATAGTTTGAGATTAACGGGGGCCGTAAAATTGCCCGTTGTAGGCCGGAT
AAGGCGTTCACGCCGCATCCGGCAAAAATTTGATTAACCGCACCTAACGGACACAACACCATGAAATCTGCCACCTCTG
CGCAAAGACCTTACCAGGAAGTCGGGGCGATGATCCGCGATCTGATCATAAAGACGCCGTACAATCCTGGGGATCCTCT
AGATTTAAGAAGGAGATATACAT**ATG**

>araEp
GGATCCCTGATGGGGCCGATAGTGTTCCTTGCCTCCAGCGCTTCAGATTATGTGAATGGTTATACCATTGCCGTGGATG
GCGGTTGGCTGGCGCGTTAATTCATTCTTCTTACTTTTATGACCCTGCCGCATGGCAGGGTTTTTTATACCTGTAGATC
ATCATAATCCATATCATGGTTATGAAATAATCCATATTAATTATCAATTAATGAACTTTATGAATTTTATCTGCTGTAA
AATTAGGTGGTTAATAATAATCTCAATAATTCAACTTAATTTGAAAATTGGAATATCCATCACATAACGACATGTCGCA
GCAATTTAATCCATATTTATGCTGTTTCCGACCTGACACCTGCGTGAGTTGTTCACGTATTTTTTCACTAT**G**TCTTACT
CTCTGCTGGCAGGAAAAAATGGTTACTATCAATACGGAATCTGCTTTAACGCCACGTTCTTTGCGGGATACGCGGCGTA
TGAATATGTTTGTTTCGGTAGCTGCTGCGCTCGAGAGATCCTCTAGATTTAAGAAGGAGATATACAT**ATG**

>ppiDp2
CTCGAGCCGCAGACCGGTAAAGAGATCACCATCGCTGCTGCTAAAGTACCGAGCTTCCGTGCAGGTAAAGCACTGAAAG
ACGCGGTAAACTAAGCGTTGTCCCCAGTGGGGATGTGACGAAGTTCAAGGGCGCATCTACTGATGTGCCTTTTTTATTT
GTATTCGGTGACTTTCTGCGTCTTGTGGGC**T**GACAATTGCCCCCGTTTCTTGTCACAATAGGCCTTTGCGCGCATCGAT
ACGTTGCGTGAGGTACACAGTCATCTACAGCGGAGTGTTGTTACACCATGATGGACAGCTTACGCACGGCTGGATCCTC
TAGATTTAAGAAGGAGATATACAT**ATG**

>aroKp1 (aroKp2 is not included)
GGATCCTCGGGCAATTATTTCGTCATGACGGAAAAGAAGATGAACGACGCGAGTTAGTGGTGTTTATCACGCCACGACT
GGTTTCCAGTGAGTAAACAGCCGTAAAAGCGGTAATGTTTTTACGCTGAACGTGTTTCATCTATTTGACGCGCGCAGGT
ATTTAGCATACAAGGAGTACC**G**ATTTGAGAGTTGGTGCTCTTCGCTGCCTGCGTTCCATGATGATGATTTATCATTCAG
GCGGCATTTTGCTGTCTTTTTTACGCTAATCTTACCCGGTGATTTATCGCCAGAGCGGTGGTAGCAAGGCAGCGCGCTT
GCAGCGACCAGATATGCAGAGGGATGGGTGATTTATTCAGTTGCCAAACCCGCTCGAGAGATCCTCTAGATTTAAGAAG
GAGATATACAT**ATG**

>aroLp
CTCGAGGGCGGACCAGATAGCCTTTCACAACGTGACCGCCAGGCCTTTGCCGCGGAGCTGGAGAAGTGGTGGCTGGAAG
TGCAACGTAGTCGTGGCTAAATGTAATTTATTATTTACACTTCATTCTTGAATATTTATTGGTATAGTAAGGGGTGT**A**T
TGAGATTTTCACTTTAAGTGGAATTTTTTCTTTACAATCGAAATTGTACTAGTTTGATGGTATGATCGCTATTCTCATG
ACACCGGCTTTCGCCGCATTGCGACCTATTGGGGAAAACCCACGATGACACAACCTCTTTTTCTGATCGGGCCTCGGGG

CTGTGGTAAAACAACGGTCGGAATGGCCCTTGCCGATTCGCTTAACCGGATCCTCTAGATTTAAGAAGGAGATATACAT
**ATG**

>aroFp

GGATCCCAACAAGGGGGCGATAAACTTTTTCATCATTTCTTTCTCCTTTTTCAAAGCATAGCGGATTGTTTTCAAAGGG
AGTGTAAATTTATCTATACAGAGGTAAGGGTTGAAAGCGCGACTAAATTGCCTGTGTAAATAAAAATGTACGAAATATG
GATTGAAAACTTTACTTTATGTGTTATCGTTACGTC**A**TCCTCGCTGAGGATCAACTATCGCAAACGAGCATAAACAGGA
TCGCCATCATGCAAAAAGACGCGCTGAATAACGTACATATTACCGACGAACAGGTTTTAATGACTCCGGAACAACTGAA
GGCCGCTTTTCCATTCTCGAGAGATCCTCTAGATTTAAGAAGGAGATATACAT**ATG**

>aroHp (aroHp1 and aroHp2 are available in this reporter)

CTCGAGTCAGATCCCGTGGATTAACAGTACCAATTATTCGGTAGAAGAGATTGCCACCAAGATCCTCGATATCATGGGC
CTTAGTCGCCGAATGTACTAGAGAACTAGTGCATTAGCTTATTTTTTTGTTATCATGCTA**A**CCACCCGGCGAGGTGTGA
CACACCTCGCACTTGAAATCAGCAGCGATTGGTTTATCGTGATGCGC**A**TCACTTCCCGGCAGTCCTGCCGTAGAAGCAA
CAAATTTCTGAGACTTGTAATGAACAGAACTGACGAACTCCGTACTGCGCGTATTGAGAGCCTGGTAACGCCCGCCGAA
CTCGCGCTACGGTATCCCGTAACGCCTGGGGATCCTCTAGATTTAAGAAGGAGATATACAT**ATG**

>pheLp

CTCGAGACAAAGGCGAAGCACGTCGTGCCGCAACATCGGTGAAAGACGCCAACTTCGTCGAAGAAGTTGAAGAAGAGTA
GTCCTTTATATTGAGTGTATCGCCAACGCGCCTTCGGGCGCGTTTTTTGTTGACAGCGTGAAAACAGTACGGGTACTGT
ACTAA**A**GTCACTTAAGGAAACAAACATGAAACACATACCGTTTTTCTTCGCATTCTTTTTTACCTTCCCCTGAATGGGA
GGATCCTCTAGATTTAAGAAGGAGATATACAT**ATG**

>secBp

GGATCCGATGGCAACGCCGCCAAGCGTGAAGAGATGATCAAACGCAGCGGTCGCACCACGGTTCCCCAGATTTTTATTG
ACGCACAGCACATTGGCGGCTGTGATGACTTGTATGCATTGGATGCACGTGGTGGACTGGATCC**C**CTGCTGAAATAACG
TGTGAACGTTGGCATTACATTGCGCAGTATTTAAGGACAACACTTAAGGGTTTTCTACACATGTCAGAACAAAACAACA
CTGAAATGACTTTCCAGATCCAACGTATTTATACCAAGGATATCTCTTTCGAAGCGCCGCTCGAGAGATCCTCTAGATT
TAAGAAGGAGATATACAT**ATG**

>serCp

CTCGAGTCATATGAAAGCGGGGGAAAAACAATTATGTCCGCGCTGTGCAAATCCAGAATGGACGAAGGCAAGTCGGGCA
AAACGGGTGACCTGACAGTAAAAACATCGGCTTTTTGCTAATAATCCGAGAGATTCTTTTGTGTGATGCAAGCCACATT
TTTGCCCTCAACGGTTTTACTCATTGCGATGTGTGTCACTGAATGATAAAACCGATAGCCACAGGAATAATGTATT**A**CC
TGTGGTCGCAATCGATTGACCGCGGGTTAATAGCAACGCAACGTGGTGAGGGGAAATGGCTCAAATCTTCAATTTTAGT
TCTGGTCCGGCAATGCTACCGGCAGAGGTGCTTAAACAGGCTCAACAGGAACTGCGCGACTGGAACGGTCTTGGGGATC
CTCTAGATTTAAGAAGGAGATATACAT**ATG**

>talAp

CTCGAGGGGATCCAATATAGCCTCTGGCAACGCAGCCTCGAGGCTGTGTTGCCAGAGGCTTGGTTGGAGAAACCTGGAT
TTTCCCTGGAACTGGAAATTCATGGAAATCAAGTGCACTTTGTTTTAACTGGTCATCCATTTGGTTGTTCCTTTCACGT
AACGTTCACAAATAAAGTGTGTGGGCAACAGCCCCTGCCCACAACGTGGCGCACATTATTACCCTGCCGGAGTCTACAG
ACTTTGAGCAAGTCCAAACTCTCACCATTAATATAATGTTTTGGTAATAATCCTATAACACTGATGTTACCTGCTTAAT
CCAGCAATACCATGCCTGTCTGCTATGCTTTTTTGATGCGTTTAGCGAAATTT**C**TCAGAAGTGTGAATTAACGCACTCA
TCTAACACTTTACTTTTCAAGGAGTATTTCCTATGAACGAGTTAGACGGCATCAAACAGTTCACCACTGTCGTGGCAGA
CAGCGGCGATATTGGATCCTCTAGATTTAAGAAGGAGATATACAT**ATG**
